# PI3K-dependant reprogramming of hexokinase isoforms controls glucose metabolism and functional responses of B lymphocytes

**DOI:** 10.1101/2024.02.29.582554

**Authors:** Brandon T Paradoski, Sen Hou, Edgard M Mejia, Folayemi Olayinka-Adefemi, Danielle Fowke, Grant M Hatch, Ayesha Saleem, Versha Banerji, Nissim Hay, Hu Zeng, Aaron J Marshall

## Abstract

B lymphocyte metabolic reprogramming is essential for B cell differentiation and mounting a healthy immune response. The PI3K signaling pathway regulates B cell metabolism, but the mechanisms involved are not well understood. Here we report that signaling via PI3Kδ can impact B cell glucose metabolism and immune functions via selective upregulation of hexokinase 2 (HK2). Three HK enzymes can catalyze the critical first step for glucose utilization and may selectively direct glucose into specific catabolic and anabolic pathways. While HK1 is constitutively expressed in B cells, HK2 is strikingly upregulated during B cell activation in a PI3Kδ-dependent manner. HK2 shows a unique distribution between mitochondrial and cytoplasmic pools that is also regulated by PI3K. Genetic deletion of HK2 significantly impairs extracellular acidification rate and glycolytic ATP production despite strong expression of HK1. B cell-specific deletion of HK2 in mice caused mild perturbations in B cell development but did not prevent generation of mature B cell subsets. HK2-deficient B cells show altered functional responses *in vitro* and evidence of metabolic adaptation to become less dependent on glucose and more dependent on glutamine. HK2-deficient B cells exhibit impaired glycolysis, altered metabolite profiles and altered flux of labeled glucose carbons into the pentose phosphate pathway. Upon immunization, HK2-deficient mice exhibit impaired generation of germinal centre B cells, plasmablasts and antibody responses. We further found that HK2 expression in primary human chronic lymphocytic leukemia (CLL) cells was associated with recent proliferation and could be reduced by PI3K inhibition. Our study identifies hexokinase 2 upregulation as a functionally important component of B cell metabolic reprogramming dependent on the PI3K pathway.

## Introduction

During an immune response, lymphocytes undergo dramatic remodeling of their cellular metabolic pathways in order to meet the demands of rapid cell proliferation and new effector functions. However, activated lymphocyte subsets exhibit variable dependence on glucose and other metabolic fuels to provide energy and building blocks needed for biosynthesis of nucleic acids and glycoproteins such as antibodies. Upon activation with antigen and costimulatory signals, B cells must rapidly transition from a quiescent state to a highly active state of rapid cell division which requires profound metabolic reprogramming. B cells reprogram their metabolism by selecting the appropriate metabolic pathways to generate energy as well as the needed biomolecules to progress through cell growth, division, differentiation, and effector function^1–3^. Resting B cells consume low amounts of macromolecules to provide energy for their basal metabolic processes and perform little glycolysis. Mitochondrial oxidative phosphorylation appears to be important for energy production, as resting B cells are sensitive to inhibition of ATP synthase using the inhibitor oligomycin and have been suggested to predominantly break down fatty acids via the process of β-oxidation. Resting B cells are maintained in their quiescent state by Glycogen Synthase Kinase 3 (GSK3) which prevents cell growth and proliferation, preserves redox levels, and inhibits glycolysis to allow for long lived B-cell quiescence^4^.

Upon activation, B cells switch to an anabolic state where ATP is more rapidly produced and utilized in building macromolecules such as proteins, nucleotides, and lipids to allow for cell growth, DNA synthesis, and cell division^5,6^. This is largely achieved by the transcription factor cMyc and kinase mTORC1 which drive new transcription and protein translation respectively to reprogram B-cell metabolism^7^, including rise in glucose transporter expression and a tenfold increase in glucose uptake^3^. Phosphatidylinositol 3-Kinases (PI3Ks) initiate a critical signaling cascade acting via PI binding proteins such as Akt, PDK1 and TAPP1/2 to control cMyc and mTOR activity and increase glucose utilization^6,8^. B cell activation via the antigen receptor alone causes B cells to initiate glycolysis and oxidative phosphorylation to support anabolic increase in cell size. Co-engagement of the inhibitory receptor FcγRIIB reduces glucose uptake, and glycolysis^5^. Antigen receptor activation without a second signal from a T cell (Eg. CD40L+IL-4) or innate immune receptor (TLR9) leads to mitochondrial swelling, increased ROS levels and dysregulated Ca^2+^ signaling, ultimately resulting in activation-induced cell death^9^. Activation via CD40L+IL-4 was found to divert glucose into the pentose phosphate pathway for nucleotides and NAPDH synthesis as well as fatty acid and cholesterol synthesis needed for cellular membrane production and cell division^5,9,10^. Taken together, the specific metabolic pathways used by acutely activated B cells are time– and signal-dependent, which coincides with the cellular changes necessary to proliferate and carry out immune functions.

Metabolic heterogeneity among different B cell subsets is not entirely understood. Anergic B cells exhibit reduced PI3K pathway activity, are metabolically quiescent and show reduced ability to produce antibodies, while chronic activation of B cells via BAFF led to enhanced metabolism which correlated with sustained PI3K pathway activity^11,12^. Selection of high affinity antigen-specific B cells during the germinal center response requires continuous cycles of cell division and affinity selection for many weeks, within a distinct hypoxic micro-environment^13,14^. We and others have found that germinal center B cells (GCB) express more GLUT1 on their cell surface and have a higher glucose uptake rate than naïve follicular B cells^4,8^. GCB receiving positive selective signals upregulate cMyc and require mTOR-dependent anabolic metabolism to undergo subsequent proliferation^15–17^. Upon differentiation into effector plasma cells (PC), B cells undergo a further metabolic transformation, devoting tremendous resources to protein synthesis and glycosylation required for high-rate Ab production^20,21^. PC take up large amounts of glucose to support energy production and synthesis of glycosylated antibodies^18,19^. Thus, GC and PC populations appear highly dependent on glucose metabolism.

Glucose enters activated B cells though glucose transporters (primarily GLUT1, 3 and 4) and is phosphorylated at the carbon six position by one of the four hexokinase (HK) enzymes to form Glucose-6-P^20^. Glucose-6-P can then be utilized by one of several important metabolic pathways such as glycolysis, oxidative phosphorylation, the pentose phosphate pathway (PPP), or the hexosamine biosynthetic pathway. Using ^13^C-glucose tracing, glucose-derived carbons can also be found interconverted into lipids, amino acids, and other sugars^1^. While mitochondrial oxidative phosphorylation has an advantage over cytosolic glycolysis in energy production, it also consumes potential sources of biosynthetic precursors and generates reactive oxygen species that can reach toxic levels. Thus, these processes must be carefully balanced within rapidly proliferating cells. Targeting glucose metabolism with drugs such as 2-deoxyglucose significantly suppressed B cell proliferation and antibody production, while B cell specific deletion of GLUT1 resulted in impaired B cell numbers or antibody production^11^.

The phosphoinositide 3-kinase (PI3K) signaling pathway is activated by antigen and costimulatory receptors and plays a central role in B cell metabolic reprograming^2,21,22^. We and others have reported that increased glucose uptake upon B cell activation is partly dependent on the PI3K pathway^8,23,24^; however, our understanding of the underlying mechanisms remains rudimentary. PI3Ks function by phosphorylating plasma membrane lipids to generate D3 phosphoinositides, which regulate the activity of over 100 PI-binding proteins^25–27^, including the extensively studied serine-threonine kinase Akt. Akt can phosphorylate and inactivate GSK3 to prevent its functions in maintaining B-cell quiescence^4^, and regulate mTOR via phosphorylation and inactivation of mTOR inhibitors PRAS40 and TSC2^28,29^. Akt1/2-deficient B cells show impaired cMyc expression and mTOR activity, reduced glucose uptake and cell growth, and mount poor GC and plasma cell responses^30^. We found that uncoupling of TAPP1/2 from the PI3K product PI(3,4)P2 leads to hyper-activation of Akt, increased glucose uptake and B cell hyperactivation *in vitro* and *in vivo*^8^. We and others have shown that PI3K can regulate expression of glucose transporter GLUT1^8,11,31–33^, suggesting that one mechanism by which PI3K promotes glucose utilization is by ensuring sufficient glucose transport capacity.

While GLUTs are bidirectional transporters facilitating diffusion across the membrane, the intracellular retention and utilization of glucose requires the action of hexokinases (HKs). HKs are enzymes that phosphorylate glucose to glucose-6-phosphate (G6P), which maintains the concentration gradient for glucose to continue entering into the cell while retaining G6P intracellularly^34^. G6P can be diverted into several distinct metabolic pathways, including glycolysis, conversion into glycogen for the storage of energy, entry into the pentose phosphate pathway (PPP) for NADPH and nucleotide production, or entry into the hexoseamine biosynthesis pathway (HBP) to glycosylate proteins^35^. There are four hexokinase protein isoforms that differ in their respective enzyme kinetics, subcellular localization, and tissue expression^36^ ^36^. While HK1 is ubiquitously expressed, other isoforms show restricted expression patterns. The HK2 isoform is expressed at very low levels in resting T lymphocytes, but is sharply elevated upon activation^37^; the expression pattern of HK isoforms in B lymphocytes has not been determined. Regulated localization of HK1 to mitochondria vs cytoplasm was recently found to impact the metabolic fate of glucose and cellular functions of macrophages^38^. In non-immune cells HK2 can shuttle between mitochondria and cytoplasm in a regulated manner^39^; however its subcellular localization in B lymphocytes remains to be determined.

Here we examined whether reprogramming of HK isoforms occurs during B cell activation, uncovering a marked upregulation of HK2 and regulated mitochondrial localization during B cell activation that both depend in part on the PI3K/mTOR pathway. We further found that HK2 has unique roles in B cell development and activation as well as generation of optimal antibody responses *in vivo*. Finally, we examine the role of HK2 in B cell metabolic re-programming during activation, providing evidence that HK2 can impact glucose utilization in glycolysis as well as direct alternate metabolic fates of glucose. Together our results strongly implicate HK2 upregulation as an important component of B cell metabolic reprogramming required for optimal glucose utilization and generation of humoral immune responses.

## Results

### Activation-induced reprograming of hexokinase isoforms is regulated by PI3Kδ activity

Previous studies have shown that inhibition of PI3K activity results in decreased glycolysis in different cell types, however its role in reprogramming glucose utilization during B cell activation is not completely understood. We examined the impact of Idelalisib, a PI3Kδ inhibitor (PI3Kδi), on activation of extracellular acidification rate (ECAR), an indirect measure of glycolysis. Primary B cells were activated overnight, with or without PI3Kδi treatment, and their resulting ECAR was measured. Our results show that PI3Kδi treatment results in a significant decrease in glycolysis as well as glycolytic reserve, while basal ECAR was not affected (**Fig 1A**). Reciprocally, primary mouse B cells expressing PI3Kδ with a gain-of-function (GOF) mutation^40^ showed markedly increased ECAR upon glucose addition (**Fig 1B**), indicating that hyper-active PI3Kδ can drive elevated glycolysis. Following uptake by glucose transporters, glucose phosphorylation by hexokinases (HKs) is a key step required to retain and utilize intracellular glucose after transport. We found that resting splenic B cells predominantly express HK1, with HK2 being expressed at very low levels; however, upon overnight stimulation, protein expression of HK2 was robustly increased while HK1 protein levels were unexpectedly found to decrease (**Fig 1C**). Interestingly, pre-treating cells with inhibitors of either PI3Kδ or all PI3K isoforms reversed this activation-dependent reprogramming of HK isoforms (**Fig 1C**). We further examined the role of two downstream targets of PI3K, Akt or mTOR, and found that inhibition of mTOR complex 1 using rapamycin or both mTOR complexes 1&2 using Sapanisertib partially blocked HK2 upregulation, whereas inhibition of Akt or mTORC2 had little effect (**Fig 1C**). Conversely, PI3Kδ-GOF B cells showed a larger increase in HK2 expression after stimulation (**Fig 1D**). Expression of HK2 in human B cells was assessed using intracellular flow cytometry analysis, with specificity of staining validated in Crispr knockout B lymphoma cells (**Fig 1E**). Primary human B cells also exhibited an increase in HK2 expression after activation, which was reversed by PI3K inhibition (**Fig 1F**). Taken together, these results indicate that B cell activation leads to PI3K-dependant increases in HK2 protein levels.

**Figure 1:**
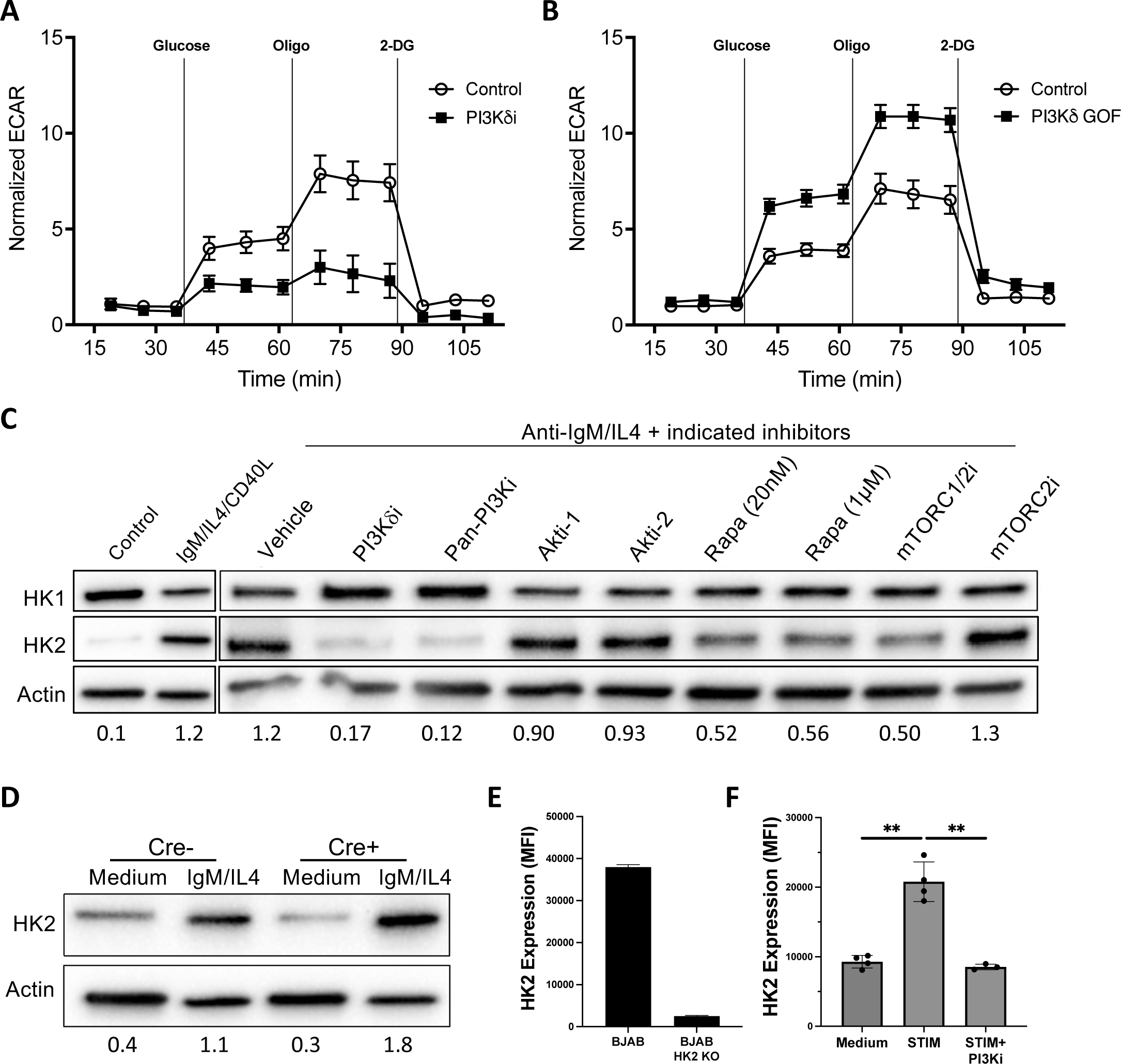
PI3K-dependant reprogramming of hexokinases during B cell activation. **A**) Splenic B cells from C57BL/6 mice were stimulated overnight with anti-CD40+IL4+anti-IgM with or without PI3Kdelta inhibitor Idelalisib (PI3K8i) and extracellular acidification rate (ECAR) profile was assessed by Seahorse assay. **B)** Splenic B cells were isolated from mice with a B cell-specific PI3K8 gain-of-function mutation (PI3K8 GOF) were stimulated overnight with anti-CD40+IL4+anti-IgM and ECAR profile assessed. **C)** Protein extracts were collected from resting or activated B cells and expression of HK1 or HK2 detected by Western blot. Impact of PI3K, Akt and mTOR inhibitors on activation-induced HK1/2 expression was determined by addition of inhibitors: PI3K8i = 10μM Idelalisib, pan-PI3Ki = 1μM Pictilisib, Akti-1 = 1μM MK-2206, Akti-2 = 1μM Iapatasertib, Rapa = rapamycin, mTORC1/2i = 50nM Sapanisertib, mTORC2i = 50nM JR-AB2-011. **D)** Expression of HK2 in PI3K8 GOF B cells. Data in A-D are representative of two to three independent experiments. **E)** Flow cytometry detection of HK2 upregulation in human B cells was validated by comparing BJAB human B lymphoma cells and BJAB cells where the HK2 gene was deleted by Crispr. **F)** Peripheral blood mononuclear cells (PBMC) were stimulated overnight with CD40L+IL4+anti-IgM with or without 1μM Pictilisib then surface stained for CD19 and intracellularly stained for HK2. HK2 staining MFI among live CD19+ cells is shown for 4 healthy donor PBMC samples. Significance was determined by paired T-test (** p < 0.005).

### HK2 exhibits distinctive subcellular localization and PI3K-regulated association with mitochondria

In addition to expression levels, the localization of HK enzymes to the mitochondria is a important factor impacting their metabolic functions. To assess mitochondrial versus cytoplasmic localization of HK isoforms in lymphocytes we utilized a human B lymphoma cell line BJAB that abundantly expresses HK1, 2 and 3. Western blot analysis of mitochondrial and cytoplasmic fractions revealed that HK1 is almost exclusively found in mitochondrial fractions, HK2 is split between mitochondrial and cytoplasmic fractions and HK3 is primarily found in cytoplasmic fractions (**Fig 2A**). We used confocal microscopy to further assess the impact of B cell activation and PI3K signaling on HK2 localization to mitochondria. Our results indicated that, under low glucose conditions, B cell activation increased HK2 localization to the mitochondria in a PI3K-dependent manner (**Fig 2B**). These results confirm differential subcellular localization of HK isoforms in B cells and suggest that PI3K signaling may also regulate subcellular localization of HK2 in the context of human B lymphoma cells.

**Figure 2:**
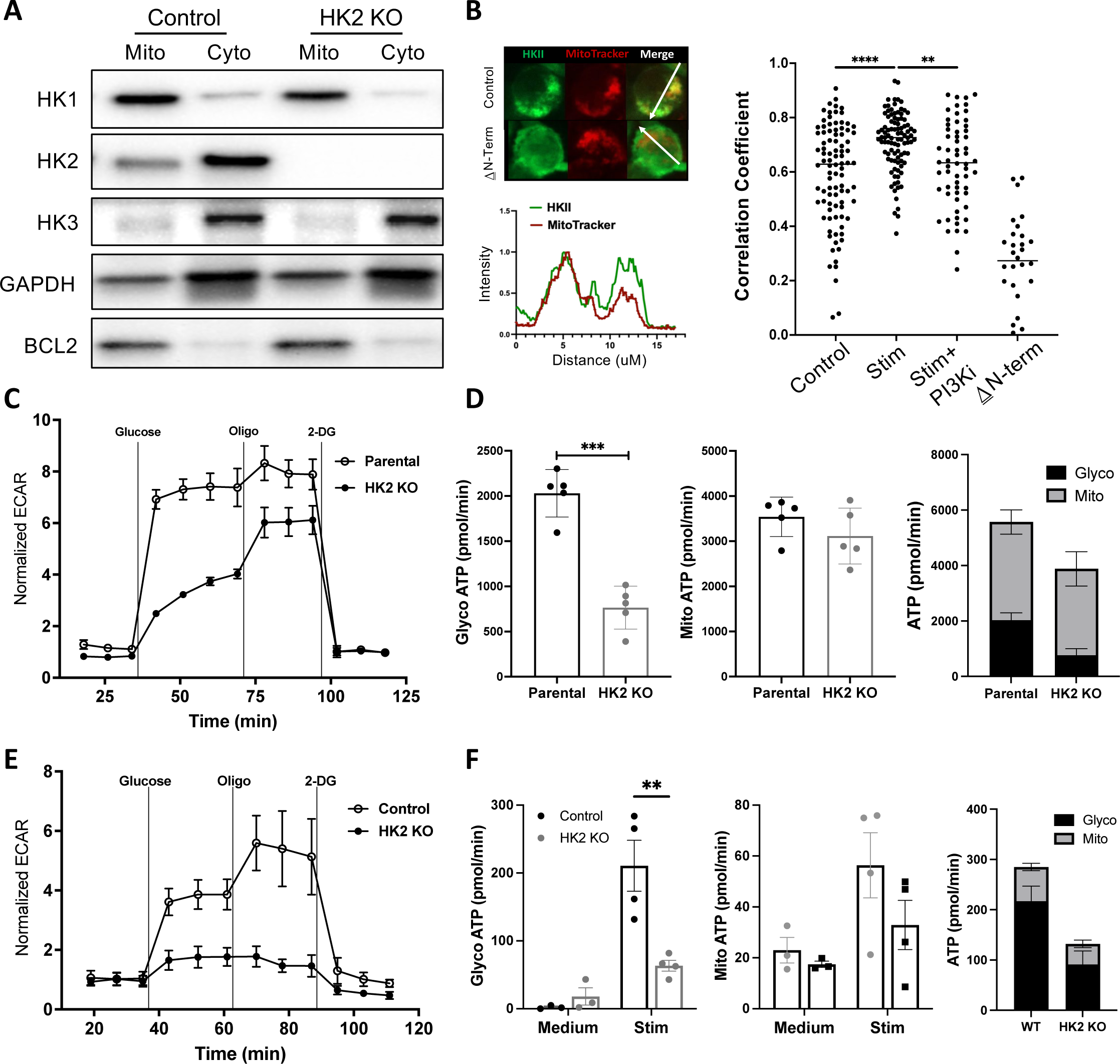
Unique subcellular localization and non-redundant function of HK2 in B cells. **A**) Mitochondrial and cytoplasmic protein extracts were prepared from parental or HK2-deficient BJAB cells and Western blotted for HK1, 2 and 3. GAPDH was used as a positive control for cytoplasmic fraction enrichment while Bcl2 was used as a positive control for mitochondrial fraction enrichment. **B)** Mitochondrial localization of HK2 in BJAB cells under various conditions was assessed by immunofluorescence staining of HK2, labeling of mitochondria with MitoTracker dye and confocal microscopy analysis. Cells were assessed under low glucose conditions (Control), after treatment with anti-CD40+IL4+anti-IgM (Stim) or pre-treatment with Pictilisib prior to stimulation (Stim+PI3Ki). HK2 KO BJAB cells re-expressing HK2 lacking the N-terminal mitochondrial localization sequence (βN-term) were used as a control. Representative images and fluorescence intensity plots are shown on the left. Graph on the right depicts Pearson’s correlation coefficients of HK2/mito staining signals for individual cells; results shown are pooled from two experiments showing similar results. **C)** Parental and HK2 deficient BJAB cells were assessed by metabolic flux assays and show significantly reduced ECAR. **D)** Metabolic flux assay results indicating that HK2 deficient BJAB cells have significantly reduced glycolytic ATP production rate. **E/F)** Splenic B cells were isolated from HK2-deficient or littermate control mice and cultured overnight in medium (control) or anti-CD40, IL-4 and anti-IgM (Stim). Glycolysis was assessed by Seahorse assays to measure ECAR in stimulated cells **(E)** or glycolytic/mitochondrial ATP production rates **(F)**. Seahorse ECAR and ATP assay results are each representative of two independent experiments. Significance was determined by unpaired T-test (* p<0.05).

### HK2 has a non-redundant function in maximizing glycolysis

To determine whether HK2 has a non-redundant functional role in B cell metabolism, we used Crispr to inactivate the HK2 gene in BJAB cells. We found that absence of HK2 protein did not affect expression or sub-cellular localization of HK1 or HK3 (**Fig 2A**). Assessment of ECAR revealed that the absence of HK2 results in a significant reduction in glycolysis (**Fig 2C**). Interestingly, parental BJAB B lymphoma cells show minimal increase in ECAR upon inhibition of oxidative phosphorylation by oligomycin, indicating that high HK2 expression is associated with high constitutive glycolytic activity and minimal glycolytic reserve; while HK2-deficient cells are able to significantly increase their ECAR levels with the addition of oligomycin (**Fig 2C**). Analysis of ATP production rate revealed that HK2-deficient cells have impaired glycolytic ATP production rate, whereas mitochondrial ATP production rate was not significantly affected (**Fig 2D**). We further examined glycolysis in HK2-deficient primary mouse B cells after overnight activation *in vitro*. A significant reduction in ECAR after addition of glucose was observed in HK2-deleted B cells compared to littermate controls (**Fig 2E**). In contrast to human B lymphoma cells, HK2-deleted mouse B cells appear to have diminished significant glycolytic reserve. Assessment of glycolytic and mitochondrial ATP production indicated a specific impairment in energy production from glycolysis (**Fig 2F**). These results suggests that HK2 has non-redundant functions to maximize glycolysis in activated B cells, but is not essential for mitochondrial ATP production.

### Impact of HK2-deficiency on B cell development

To investigate the roles of HK2 in normal B cell development and function, we utilized mice with B cell-specific inactivation of the HK2 gene. We confirmed HK2 deficiency in splenic B cells at the mRNA and protein levels, while HK1 and HK3 expression remained intact (**Fig S1**). We then performed intracellular flow cytometry analysis in various mouse tissues to determine HK2 protein levels during B cell development and activation (flow gating illustrated in **Fig S2**). The specificity of rabbit monoclonal anti-HK2 reagent used was validated using HK2-deficient activated mouse splenic B cells (**Fig 3A**). In bone marrow, HK2 expression is low in all B cell subsets with the exception of pro-B cells which express significantly more HK2 than other B lineage subsets (**Fig 3B**). In spleen, mature follicular B cells express the lowest levels of HK2, with marginal zone and B1 populations expressing significantly higher levels (**Fig 3C**). In peritoneal wash, B1 cells were also observed to express higher levels of HK2 than B2 cells (**Fig 3D**). Interestingly basal populations of CD86+ B cells and germinal center cells in spleen, mesenteric lymph nodes and Peyers patches showed substantially higher expression of HK2 than resting B cells in these same tissues (**Fig S3A**). These results are relatively consistent with levels of HK2 mRNA transcripts in B cell subsets as reported in the Immgen database (**Fig S3B**).

**Figure 3:**
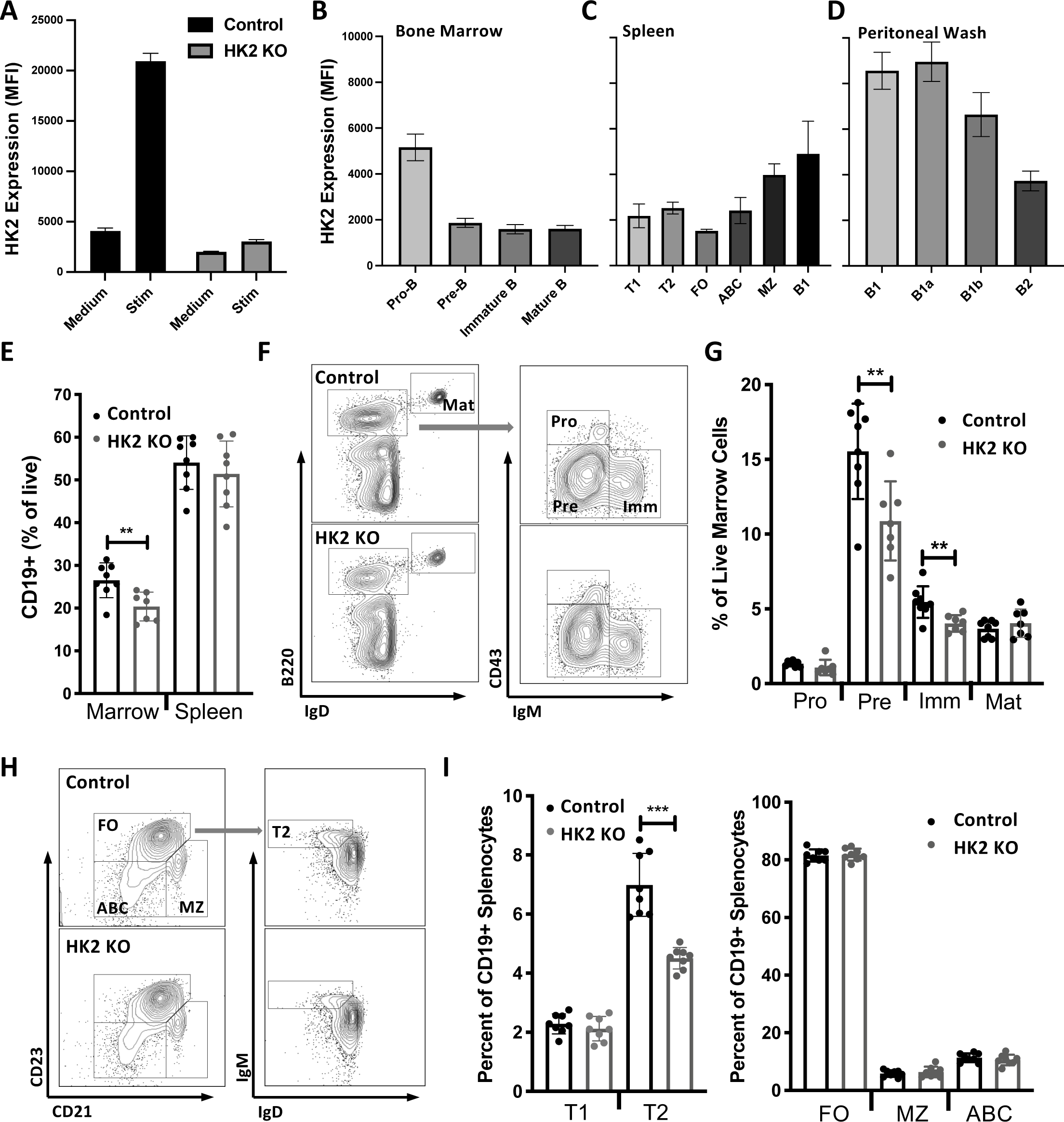
Roles of HK2 in B cell development. **A**) Validation of HK2 intracellular staining specificity using activated HK2 KO B cells. (**B-D)** Single cell suspensions from the indicated mouse tissues were surface stained to identify B cell sub-populations and intracellularly stained to assess HK2 expression. Background-subtracted HK2 mean fluorescence intensities (MFIs) were determined for each cell population. T1/T2=transitional 1/2; FO=follicular; ABC=age-associated B cell; MZ=marginal zone. Population gatings are described in Supplementary Figure 2. Results show mean and standard deviation of 5 mice per group and are representative of 2-3 experiments per tissue. **E)** Frequencies of CD19+ B cells in spleen and bone marrow. **F)** Representative flow cytometry plots illustrating gating in bone marrow. **G)** Frequencies of B cell and precursor subsets in bone marrow. **H)** Representative flow cytometry plots illustrating gating of CD19+ B cell subsets in spleen. **I)** Frequencies of immature and mature B cells subsets in spleen. Results in E-I are pooled from two experiments, totalling 8 mice per genotype. Significance was determined by T-test (** p<0.005; *** p<0.001).

While HK2 deficient mice exhibited normal frequencies of CD19+ B cells in spleen, peritoneal wash, mesenteric lymph nodes and Peyer Patch; however, a small but significant reduction in B lineage cells within the bone marrow was noted (**Fig 3E; Fig S4A**). Examination of B cell subsets in bone marrow revealed that the reduced frequency of B lineage cells was due to significantly reduced pre-B and immature B cells (**Fig 3F/G**). While mature B cell subsets were present at normal frequencies in spleen, a reduced frequency of transitional 2 B cells was observed in HK2-deficient mice (**Fig 3H/I**). Frequencies of spontaneous germinal center B cells in mesenteric lymph node, Peyer’s patch and spleen were within normal ranges (**Fig S4B**) and B1a, B1b and B2 cell subsets in peritoneal cavity were also not affected (**Fig S4C**). Basal serum levels of all antibody isotypes tested were within normal ranges, except IgM which showed a small decrease in HK2-deficient mice (**Fig S4D**). These results indicate that HK2 deficiency perturbs B cell development, but does not block generation of mature B cell subsets.

### HK2 deficiency impacts B cell functional responses in vitro

B cells isolated from HK2-deleted B cells or littermate controls were activated overnight and their functional responses assessed. We found that HK2-deficient B cells show slightly reduced proliferation in response to LPS or LPS+IL4, while proliferation in response to anti-CD40+IL-4 was not significantly affected (**Fig 4A**). HK2-deficient B cells showed markedly impaired IgG1 secretion with either LPS+IL4 or CD40+IL4 stimulation (**Fig 4B**). To examine whether HK2 deficiency renders B cells more sensitive to glycolysis inhibition, we assessed the effect of glycolysis inhibitor 2-deoxyglucose (2DG). We unexpectedly found that HK2 deficient B cells were partially resistant to the inhibitory effects of 2DG on proliferation (**Fig 4C**) and production of cytokines (**Fig 4D**). This suggested that HK2-deficient B cells may undergo compensatory adaption of their metabolic network to be less dependent on glycolysis. Since glutamine is an additional important source of energy and biosynthetic precursors in activated lymphocytes, we examined the sensitivity of HK2-deficient B cells to the glutaminase inhibitor CB-839 (Telaglenastat). Remarkably, HK2-deficient B cells are significantly more sensitive to glutaminase inhibition, showing greater reductions in proliferation, IgM secretion and IgG1 secretion in the presence of CB-839 (**Fig 4E**). Together these results suggest that HK2 normally plays a significant role in B activation, but in the face of congenital HK2 deficiency, B cells metabolically adapt to reduce their dependence on glucose in part by increased utilization of glutamine as an alternative fuel source.

**Figure 4:**
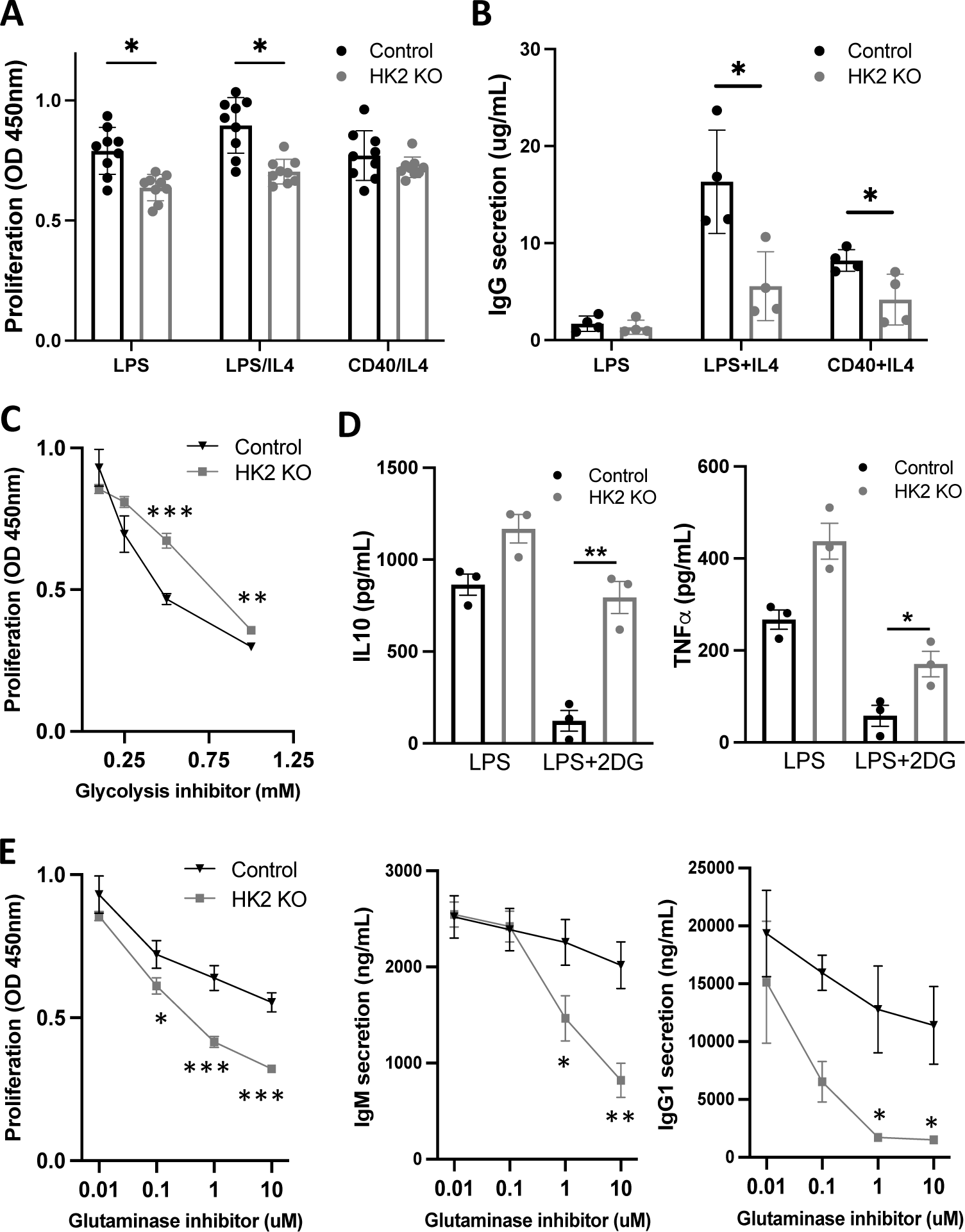
HK2 deficiency impacts in vitro B cell functional responses. Splenic B cells were isolated from HK2-inactivated or littermate control mice and cultured in the presence of the indicated activation stimuli. **A)** Proliferation was assessed at day 4 of culture by CCK-8 assay. Results in are pooled from two experiments, totalling 8 mice per genotype. **B)** After 6 days of culture, supernatants were collected to measure secreted IgG1 antibodies by ELISA assay. The experiment shown (4 mice per genotype) is representative of 3 experiments with similar results. **C)** Cells were stimulated with LPS+IL4 in the presence of the indicated concentrations of glycolysis inhibitor 2-deoxyglucose (2DG) and proliferation assessed. **D)** Cells were stimulated with LPS in the presence of 0.5mM 2DG and the indicated cytokines were assessed by Mesoscale assay. **E)** Cells stimulated with LPS+IL4 in the presence of the indicated concentrations glutaminase inhibitor CB-839 and proliferation or antibody secretion were assessed. Results in C and D are representative of 3 independent experiments; cytokine results are from a single experiment (3 mice per genotype). Significance was determined by T-test (*p<0.05; **p<0.005; ***p<0.001).

### Impact of HK2 deficiency on B cell metabolite profiles

To determine how metabolic pathways are altered in HK2-deficient B cells, a panel of 84 metabolites in central carbon metabolism were measured by mass spectrometry, comparing splenic B cells from control or HK2-deficient mice cultured overnight in medium or stimulation cocktail (**Fig 5A; Supp Table 1**). Both lactic acid and TCA cycle intermediates were reduced in activated HK2-deficient cells (**Fig 5A**). As expected, glucose-6-P/glucose ratios were decreased in HK2-deficient cells, while glycolytic intermediate fructose-6-P was only modestly reduced relative to glucose (**Fig 5B**). Pentose phosphate pathway (PPP) intermediate 6-phosphogluconate as well as glycogen synthesis intermediate glucose-1-P were both reduced relative to glucose (**Fig 5B**). Notably, HK2-deficient B cells exhibit substantial (1.5-4 fold) reductions in most nucleotides measured under resting conditions, however this deficiency was largely overcome upon stimulation (**Fig 5C** left versus middle panel). Treatment of cells with 2DG during stimulation led to a reversed trend, with HK2 KO cells showing higher levels of nucleotide metabolites under this condition (**Fig 5C**). After incubation of cells with ^13^C glucose, we assessed the extent of ^13^C labelling in a number of downstream metabolites (**Fig 5D**; **Fig S5**). Reduced labeling of 6-phosphogluconate was observed, potentially indicating impaired entry of glucose into the pentose phosphate pathway (**Fig 5D**). We also observed reduced 13C labeling of 2-phosphoglycerate (**Fig 5D**), suggesting an increased proportion of unlabeled metabolites such as amino acids feeding into the 2-PG pool. Subtle differences in labeling of TCA metabolites were observed in stimulated HK2 KO cells (**Fig S5**); notably, all TCA metabolites and direct TCA derivatives measured show approximately two-fold increases in M+1 labelling, and citrate, isocitrate, and succinate also showed corresponding reductions in M+2 labelling, potentially reflecting a reduced proportion of glucose-derived carbons entering into the TCA. Overall, these findings suggest that HK2-deficient B-cells have defects across several anabolic and catabolic pathways but can overcome some of these deficiencies with a strong activation stimulus, likely via increased utilization of other sugars or amino acids.

**Figure 5:**
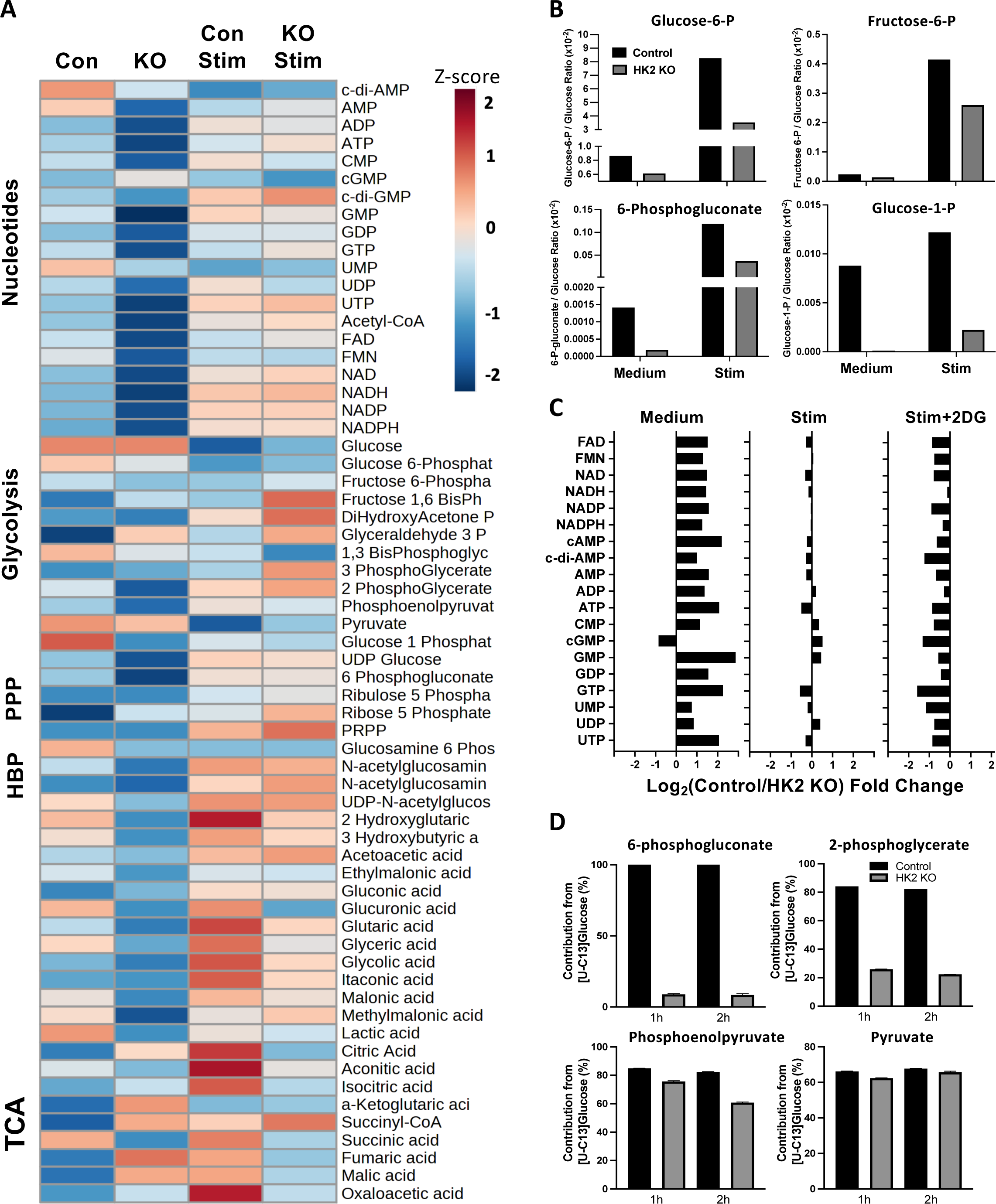
Impact of HK2 deficiency on metabolite profiles. Splenic B-cells were cultured overnight in RPMI media only or with F(ab’)2 anti-IgM, CD40L and IL4 (STIM), snap frozen and metabolites measured by mass spectrometry. Metabolite levels were normalized to total protein content in each sample. **A)** Heat map illustrating relative differences in metabolite levels. **B)** Calculated ratios of glucose-6-P, fructose-6-P, 6-phosphogluconate or glucose-1-P to glucose under resting or activated conditions. **C)** Impact of HK2 deficiency on nucleotide metabolites, in resting, stimulated and 2DG treated conditions. Data are expressed as log transformed fold change (positive numbers indicate reduced levels in HK2 KO and negative numbers indicate elevated levels in HK2 KO relative to controls). **D)** Cells were stimulated overnight, washed in glucose-free medium and incubated for 1 or 2 hours with ^13^C-glucose to allow incorporation of ^13^C into downstream metabolites. ^13^C-labelled and unlabelled metabolites were measured by mass spectrometry and percent labelling of each metabolite is shown.

### Impact of HK2 deficiency on response to immunization

To further assess HK2 expression and function *in vivo*, mice were immunized with sheep red blood cells (SRBC) to generate strong germinal center and plasmablast responses. Germinal center B cells and plasmablasts in the spleen displayed very high HK2 expression peaking early in the response and maintained up until day 8 (**Fig 6A**). Flow cytometry assessment at day 8 post-immunization revealed significant alterations in activated cell populations, including reductions in CD86+ cells and germinal center B cells (**Fig 6B**). Plasmablast frequencies were also reduced in the spleen and bone marrow of SRBC-immunized HK2-deleted mice (**Fig 6C**). SRBC-binding IgM and IgG antibodies were significantly reduced post-immunization (**Fig 6D**). Together these results indicate that HK2 deficiency leads to significant impairments in B cell functional responses to SRBC immunization.

**Figure 6:**
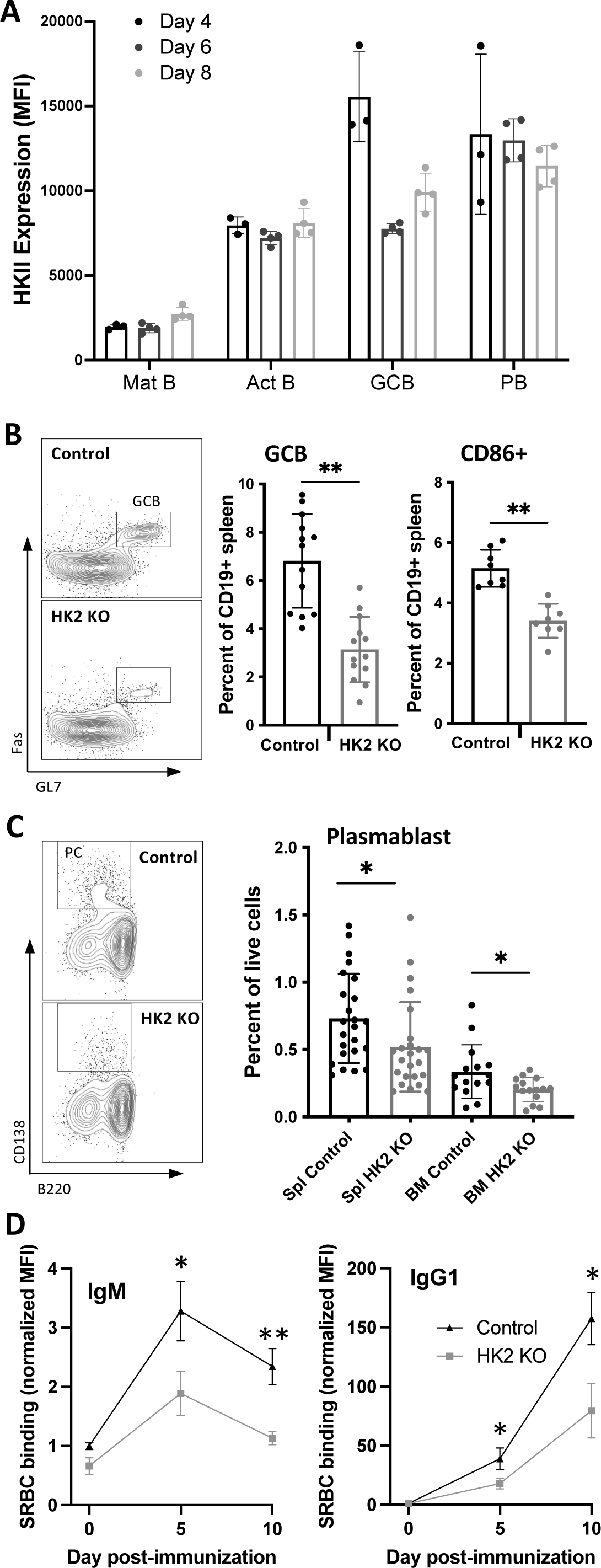
Effect of B cell-specific HK2-deficiency on responses to immunization. HK2-deficient or littermate control mice were immunized with sheep red blood cells (SRBC) and assessed by flow cytometry. **A)** HK2 expression in activated B cell populations generated at the indicated time points after sheep red blood cell immunization. **B)** Representative flow cytometry plots illustrating gating of germinal center B cell populations in spleen and graphs showing frequencies of germinal centre B cells and CD86+ B cells pooled from 2-3 experiments. **C)** Representative flow cytometry plots illustrating gating of plasma cell populations in spleen and bone marrow and graph showing cell frequencies pooled from 5 experiments. **D)** Generation of sheep red blood cell (SRBC)-binding IgM and IgG antibodies. SRBC were incubated with serum from the indicated days post-immunization, and bound antibodies were detected by immunofluorescence staining and flow cytometry. Results are representative of 4 similar experiments. Significance was determined by T-test (*p<0.05; **p<0.005).

### HK2 expression in malignant human B cells

B cell malignancies such as chronic lymphocytic leukemia (CLL) are known to be driven by chronic PI3K signaling^41^ and exhibit metabolic reprogramming^42,43^. We thus assessed HK2 protein expression in a cohort of CLL patients using flow cytometry. Directly *ex vivo*, CLL cells show a trend of reduced HK2 expression compared to healthy control B cells (**Fig 7A**), potentially reflecting the predominantly quiescent phenotype of CLL cells in blood. We determined association of CLL HK2 expression with various prognostic markers and found that it is significantly associated with ZAP70 expression (**Fig 7B**). While a trend of higher HK2 expression was observed in IgVH unmutated patients, it was not associated with lymphocyte counts or B2M levels (**Fig S6**). Strikingly, higher HK2 expression was consistently seen within the proliferative fraction of CLL cells, characterized by low levels of surface CXCR4 and high expression of CD5 (**Fig 7C**)^44,45^. Consistent with this, HK2 expression was inversely correlated with surface CXCR4 level (**Fig 7D**). Finally, we cultured CLL cells overnight with an activation cocktail, with or without PI3K inhibitor, and found significant reduction in HK2 expression upon PI3K inhibitor treatment (**Fig 7E**). These results suggest that HK2 expression may be upregulated by activation signals within lymphoid tissues and subsequently diminish in long-lived quiescent CLL cells in the blood.

**Figure 7:**
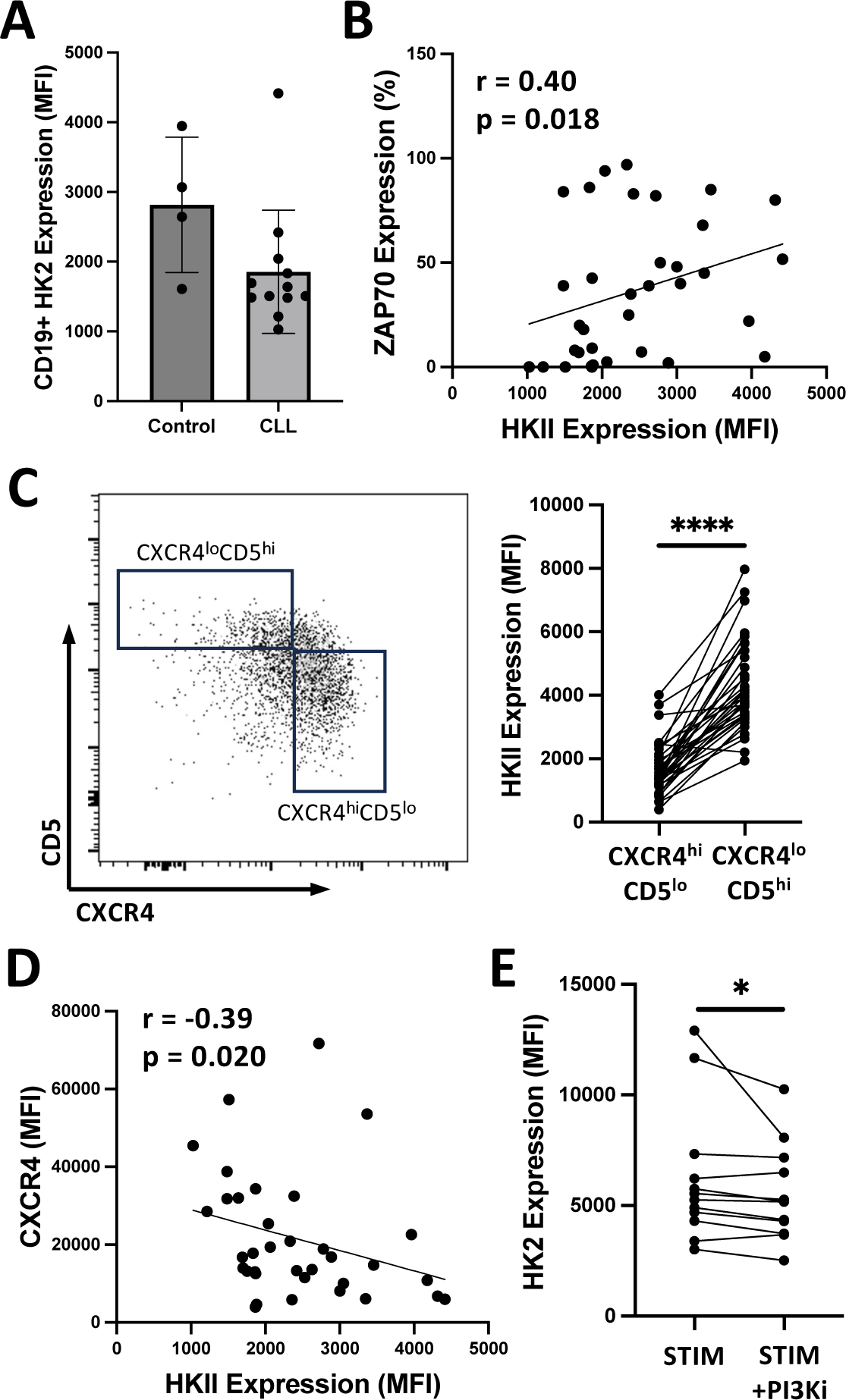
Expression of HK2 in B cell leukemia. PBMC samples from chronic lymphocytic leukemia patients or healthy controls were assessed for HK2 expression by flow cytometry. **A)** HK2 mean fluorescence intensity (MFI) within gated CD19+ cells. Each dot represents an individual patient or healthy control. **B)** Spearman’s correlation of HK2 MFI with percent ZAP70 expression among CLL cells (determined by clinical lab testing). **C**) Proliferative versus quiescent CLL cell populations were gated based on CXCR4 and CD5 expression (left panel). HK2 expression in individual patient’s proliferative and quiescent cell fractions are connected by lines (right panel). Significance was determined by Wilcoxon test. **D)** Pearson’s correlation of HK2 MFI with CXCR4 expression among CLL cells**. E)** CLL cells were cultured overnight with CD40L+IL4+anti-IgM (STIM) alone or with pan-PI3K inhibitor (1μM Pictilisib) prior to assessment of HK2 expression. Significance was determined by Wilcoxon test.

## Discussion

It is well established that B cell activation increases glycolysis and this has largely been attributed to elevation of the glucose transporter Glut1 and increased glucose uptake. Our study provides multiple lines of evidence indicating that the increase in glycolysis upon B cell activation is also associated with selectively increased expression and activity of hexokinase 2. As the first enzyme to act upon glucose entering the cell, hexokinases play a critical role, as conversion to glucose 6-phosphate prevents export and irreversibly directs glucose to multiple downstream anabolic and catabolic pathways. Although B cells constitutively express hexokinase 1, our results indicate that activated B cells predominantly express HK2 and are functionally dependent on this enzyme to maximally activate glucose-dependant metabolism. Our study indicates that signaling through the PI3K pathway, and PI3Kδ specifically, is involved in reprogramming of HK isoforms, in addition to the previously reported role of PI3K in regulating Glut1 expression. The functional importance of HK2 is underlined by our findings that HK2-deficient B cells have impaired responses to stimulation *in vitro* and *in vivo*, despite showing evidence of metabolic reprogramming to reduce their dependence on glucose.

Substantial literature has implicated the PI3K signaling pathway in metabolic reprogramming in multiple cell types. In many cases, the evidence is based on pan-PI3K inhibitors or gain-of-function approaches of limited specificity. Our data specifically implicate PI3Kδ in activation-induced increase in glycolysis and elevation of HK2 expression. The mechanisms linking PI3K to glycolysis have been proposed to include Akt-dependant upregulation of Glut1, primarily based on experiments over-expressing membrane-targeted Akt^33,46–48^. The PI3K pathway is also closely intertwined with mTOR activation^49^, which is a critical regulator of cellular metabolism^50,51^. Our results suggest that mTORC1 does play a role in HK2 upregulation in B cells, whereas Akt kinase activity is surprisingly not required. We have previously validated the impact of the inhibitors used on Akt phosphorylation and activity^8^, and thus conclude that Akt kinase activity does not play an essential non-redundant role under the activation conditions used here. PI3K and Akt can also play a role in upregulation of c-myc which is a key driver of glycolytic gene expression, including HK2^52–54^, thus c-myc may also contribute to the PI3K-dependant upregulation we observe here.

While all hexokinase isoforms catalyze the same reaction, they differ in expression pattern, structure, enzyme kinetics and subcellular localization. Unlike other isoforms, HK2 has two functional catalytic domains but a relatively low affinity for glucose^55^, and is thus expected to be most important when cells are highly active in taking up glucose. While all hexokinase isoforms have the same catalytic activity, they differ in expression pattern, structure and subcellular localization. HK2 has two functional catalytic domains and is thought to be the most active kinase in the family. While HK1 is mainly localized to mitochondria and HK3 is mainly cytoplasmic, HK2 is found in both mitochondria and cytoplasm. In the present study we found evidence that the PI3K pathway can impact mitochondrial localization of HK2, suggesting an additional layer of regulation beyond protein expression levels. This could be due in part to phosphorylation of HK2 by Akt which was reported to enhance HK2 mitochondrial association^56^; however other kinases such as PIM2 may also serve this function^57^ and constitutively active Akt does not significantly increase mitochondrial HK activity^47^. Several studies have found that dissociating HK2 from mitochondria can trigger mitochondrial dysfunction, disrupted calcium homeostasis and cell death in some contexts^58^, indicating the functional importance of HK2 mitochondrial association. A recent study found that HK1 can also undergo regulated translocation between mitochondria and cytoplasm in macrophages, and found that mitochondrial localization of HK1 was functionally important for macrophage activation^38^. Together, the current evidence indicates that PI3K and other signaling pathways can reprogram both HK isoform expression and subcellular localization, likely impacting the utilization of glucose in different downstream metabolic pathways.

Our assessment of HK2 protein expression in B cell subsets indicates it could potentially have roles in B cell development including at the pro-B and transitional B cell stages. While HK2 is not essential for B cell development, HK2 KO mice exhibit mild impairments in key developmental transitions known to be dependent on BCR signaling and the PI3K pathway. We see significant reductions in bone marrow pre-B cells and spleen transitional 2 cells, known to depend on pre-BCR or BCR signaling respectively. Although B1 and marginal zone B cells showed higher levels of HK2 than follicular B cells, all of these mature B cell populations were present at normal frequencies in HK2-deficient mice. It’s possible that metabolic plasticity allows developing B cells to proceed through checkpoints and adapt their metabolic pathways to compensate for the absence of HK2. We did not observe substantial compensatory increases in HK1 or HK3 expression, although a trend of marginally increased HK3 was noted in splenic B cells. Together our results are more consistent with metabolic adaptations that reduce B cell dependence of glucose and potentially increase their dependence on other sources of energy and metabolic building blocks such as amino acids or other sugars.

We find that deletion of HK2 in human lymphoma cells or primary mouse B cells substantially reduced extracellular acidification rate upon glucose addition, despite continued expression of HK1 and HK3. This indicates that HK2 plays an essential non-redundant role in maximizing glycolytic capacity that is not easily compensated. Surprisingly, this did not alter the growth rate of HK2 deficient BJAB cells (data not shown). HK2-deficient mouse B cells similarly show relatively normal proliferative responses. The mild impairment of proliferative responses to LPS we observed using an MTT-based assay is likely due to slightly reduced cell survival of HK2 KO cells in these cultures (data not shown). Strikingly, proliferation of HK2 KO B cells was found to be less affected by glycolysis inhibition and more sensitive to glutaminase inhibition, indicating that these cells have adapted to become less dependent on glycolysis and instead rely more on glutaminolysis. This adaptation illustrates an unexpected level of metabolic plasticity in B cell responses, which may somewhat obscure the normal roles of HK2 when cells are cultured in media with abundant glucose and glutamine.

HK2-deficiency significantly impacted metabolite profiles of splenic B cells, again demonstrating that this enzyme has metabolic functions which cannot be fully compensated by HK1/3. As metabolite levels are influenced both by rates of production and consumption, the interpretation of these data are complex. For example glucose levels are found to substantially decrease upon B cell activation, despite known increases in glucose uptake, presumably reflecting more rapid consumption by phosphorylation into glucose 6-phosphate and utilization in downstream pathways. Consistent with reduction in ECAR, TCA metabolites and lactic acid were reduced in activated HK2 KO cells compared to controls. However entry of glucose into glycolysis appeared only marginally impaired, based on glucose/fructose-6-P ratios and reduced ^13^C-glucose labeling of some glycolysis intermediates. While HK2 KO cells were found to have significantly decreased levels of nucleotides, intermediates in utilization of glucose for nucleotide synthesis, such as ribose-5-P and PRPP, were not decreased; this might reflect decreased consumption of these metabolites due to impaired pentose phosphate pathway activity. Consistent with this interpretation, HK2-deficient cells show reduced 6-phosphogluconate/glucose ratios as well as reduced incorporation of ^13^C-glucose into 6-phosphogluconate; this suggests a substantial impairment in the first step of glucose entry into the PPP. This conclusion is consistent with a recent study showing that glucose largely flows into the PPP in activated B cells and this pathway is impaired in Glut1-deficient cells^59^. Glucose conversion into glucose-1-P, the first step in glycogen synthesis, also appeared to be severely impacted by HK2-deficiency based on low glucose-1-P/glucose ratios. While little is known about the role of glycogen storage in lymphocytes, they do express glycogen synthase, contain measurable stores of glycogen and functional insulin receptor^60,61^. Together our results indicate that HK2-deficiency substantially impairs glucose utilization, particularly for anabolic pathways, and may impact glucose storage.

Upon immunization with sheep red blood cells, we found that B cell-specific HK2-deficiency caused a significant reduction in both germinal center and plasma cells. This is consistent with the observed elevation in HK2 protein levels in germinal center and plasma cells and suggests these cells may be particularly dependent on HK2 activity. Several studies have indicated that the germinal centre is a hypoxic environment, and consequently GC B cells increase activation of glycolysis pathways^14,24^; however, other studies have highlighted the importance of oxidative phosphorylation in GC B cells^14,62^. Interestingly, a recent study examining lactate dehydrogenase-deficient B cell responses concluded that aerobic glycolysis is most important in early (day 4) pre-GC GL7+ cells^63^. As we noted that HK2 expression within GC phenotype cells peaked at day 4 post-immunization, its likely that HK2 helps drive glycolysis at this stage. Plasma cells are known to exhibit high glucose utilization, with a large fraction being utilized for protein glycosylation^18^. Thus, it’s possible that the defective antibody responses we observe relate to impaired plasma cell generation, survival and/or reduced ability to secrete antibody. Notably, a recent study found that the requirement for glucose uptake to sustain plasma cell responses in vitro could be compensated by addition of other hexoses (galactose and mannose)^59^, thus glucose may not be absolutely required if sufficient supply of other sugars is available to sustain high level production of glycosylated antibodies.

While human B lymphoma lines such as BJAB constitutively express very high levels of HK2, primary CLL cells in blood did not exhibit elevated HK2 levels compared to normal human B cells. This is consistent with studies showing that the bulk of blood CLL cells are quiescent and rely mainly on mitochondrial lipid oxidation^64^. However, we found evidence that the recently proliferated fraction of CLL has markedly elevated levels of HK2, suggesting that actively dividing CLL cells may upregulate HK2 and enhance glucose utilization within lymph node proliferation centers. This upregulation may occur in response to activation signals present in this location that trigger the PI3K pathway^65^.

Together our results suggest that acute inhibition of hexokinase 2 activity could be applied to selectively inhibit germinal center and plasma cell responses as well as proliferation of malignant B cells. Indeed 2DG treatment in vivo has been found to disrupt chronic germinal centers and reduce autoantibody generation *in vivo*^8,66^, which may be in part attributable to direct effects on GCB and PC. However, given the metabolic plasticity observed in this study, combined inhibition of hexokinase 2 and other compensatory pathways may be more effective.

## Materials and Methods

### Cell culture

The human B lymphoma cell line BJAB was seeded at a concentration of 5 x 10^5^ cells/mL in RPMI (Hyclone Laboratories, #SH30027.01) containing 10% FBS (ThermoFisher, #12483-020) and 1% pen-strep (GE Healthcare Life Sciences, #15070-063) and incubated at 37°C and 5% CO_2_. Peripheral blood mononuclear cells from healthy donors or CLL patients were isolated and cultured in the same medium at 2×10^6^ per mL. Murine splenic B cells were isolated by negative selection using the EasySep Mouse B Cell Isolation Kit (STEMCELL Technologies, #19854A) as specified by the manufacturer’s protocol. B cells were seeded at a concentration of 2×10^6^ cells/mL in RPMI containing 10% FBS, 50µM 2-mercaptoethanol and 1% pen-strep and incubated at 37°C and 5% CO_2_. Where indicated, cells were stimulated with 2µg/mL of anti-CD40 (BD Pharmingen, #553721), 10ng/mL of IL4 (Peprotech, #214-14-20UG) and 10µg/mL of anti-IgM (Jackson ImmunoResearch Laboratories, #115-006-020). Human B cell stimulation used 10ng/mL Goat F(ab’)_2_ Anti-human IgM (Southern Biotech #2022-01), 2ug/mL of recombinant human sCD40 (invitrogen PHP0025), and 10ng/mL of recombinant human IL-4 (R&D systems, #204-IL/CF). Inhibitors used were the PI3Kdelta inhibitor Idelalisib, pan-PI3K inhibitor Pictilisib, Akt inhibitors Iapatasertib or MK-2206, mTOR inhibitors rapamycin, Sapanisertib or JR-AB2-01 or glutaminase inhibitor Telaglenastat (all from Selleck Chemicals). 2-deoxyglucose was purchased from Sigma-Aldrich (#D3179).

### Crispr deletion of HK2

The procedure was carried out according to a protocol generously provided by Dr. Marco Cavallari from the University of Freiburg, with minor modifications. Briefly, BJAB cells were seeded at a concentration of 5 x 10^5^ cells/mL in in RPMI containing 10% FBS and 1% pen-strep and incubated at 37°C and 5% CO_2_ 24 hours prior to the experiment. Predesigned guide RNA (crRNA:tracrRNA complex) specific for HK2 was purchased from Integrated DNA Technologies (Alt-R system) and assembled into a ribonucleoprotein complex with Cas9 according to the manufacturer’s protocol. BJAB cells were electroporated with the RNP complex using the Neon transfection system (ThermoFisher) as follows. Cells (1×10^6^) were resuspended in 9uL of buffer R, mixed with 1 uL RNP complex and 2uL of 10.8 uM Alt-R Electroporation Enhancer, and electroporated at 1350V, 40ms, 1 pulse using a 10uL Neon pipette tip. Cells were then transferred to 1mL of recovery medium in a 24 well plate (RPMI+20%FCS+Glutamax+10mM HEPES). After 3 days, cells were cloned by limiting dilution and later screened by Western blotting. Approximately 50% of clones exhibited loss of HK2 protein.

### Mice

HK2-flox mice were generously provided by Dr Nissim Hay (University of Illinois)^37^ and crossed with MB1-Cre (Jackson Labs) to generate B cell specific deletion. Genotyping and monitoring for germline deletion was done as described^67^. Mice bearing the PI3KD^E1021K^ mutation in B cells^40^ were generously provided by Dr. David Rawlings (University of Washington). All animals were housed in a specific pathogen-free facility at the University of Manitoba, according to the Canadian Council on Animal Care guidelines and used for experiments between 8-12 weeks of age.

### RT-qPCR

Splenic B cells were stimulated as indicated and RNA was isolated using RNeasy Plus Mini Kit (Qiagen). RT-PCR was carried out as described^8^ using the following primers: Fwd HK1 – CGCAGCTCCTGGCCTATTAC; Rev HK1 – GAGCCGCATGGCATAGAGAT; Fwd HK2: AGTGGAAGGCAGAGACGTTG; Rev HK2 – CAGTGCGAATGTCGTTGAGC; Fwd HK3 – TTGGGGTGCCTCATATTGCC; Rev HK3 – CTGTGCCCTTGTCACCTTGA

### Western blots

Western blots were carried out essentially as described with the following modifications. SDS-PAGE was performed using 10 μg of NP-40 extract protein mixed with an equal volume of 2X Laemmli buffer (BioRad cat#: 1610737) supplemented with 5% 2-mercaptoethanol (without boiling). Mitochondrial and cytoplasmic protein was isolated using the mitochondrial/cytoplasmic fractionation kit from Millipore Sigma (cat.#: MIT1000). Primary antibodies used include the following: HK1 (Cell Signalling Technology; cat.# 2024S), HK2 (Cell Signalling Technology; cat.#: 2867S), HK3 (ThermoFisher Scientific, cat.#: PA5-84306), β-actin (Cell Signalling Technology; cat.#: 4970S), Bcl2 (Millipore Sigma; cat.#: 05-729-25UG), GAPDH (Millipore Sigma; cat.#: CS204254). Blots were visualized using the Clarity™ Western ECL Substrate kit (Bio Rad, cat.#: 170-5061) and imaging on a ChemiDoc™ MP imaging system. The relative band intensities were determined using ImageJ software.

### Confocal microscopy analysis of mitochondrial localization

BJAB cells were resuspended in Dulbecco’s-PBS (Gibco 14040-133) containing 10mM HEPES (Gibco 15630-106) and plated at 5×10^5^ cells/mL for two hours with the indicated stimuli/inhibitors. Cells were then stained with fixable mitochondrial dye and rabbit anti-HK2 as follows. Cells were resuspended in PBS+2% FBS buffer containing 200nM of MitoTracker Deep Red FM dye (M22426 Invitrogen) and incubated for 30min at 37 degrees in 5%CO2. After washing with PBS, cells were fixed with 2% paraformaldehyde, permeabilized with 0.5% saponin, stained for 3 hours with a rabbit monoclonal anti-HK2 (Abcam #epr209847) at a 1:100 dilution, and finally stained with secondary AlexaFluor488-labeled goat anti-rabbit IgG (Invitrogen #A11034) at dilution of 1:200 for 30 minutes. Stained cells were then centrifuged onto microscope slides (Fisher #12-550-123) using a Shandon Cytospin 2 (2min x 1200 RPM) and mounted using ProLong™ Gold Antifade Mountant with DAPI (Thermo #P36935). Slides were imaged at 63X magnification using a Zeiss AxioObserver CSU-X1M 5000 spinning disc confocal microscope. At least 30 cells per group per experiment were imaged and Pearson’s Co-Localization Coefficients were determined using the Zeiss software.

### Metabolic flux assays

Extracellular acidification rate (ECAR) assays were run using a Seahorse XF24 instrument essentially as described^8^. ATP production rates were determined using the Seahorse XF Real-Time ATP Rate Assay Kit (Agilent 103591-100), according to the manufacturer’s protocol. Primary mouse B cells were resuspended in the supplemented XF DMEM media to a concentration of 4×10^6^ cells/mL while BJAB cells were resuspended at a concentration of 2×10^6^ cells/mL and applied to assay plates coated with Poly-D-Lysine (Gibco A38904-01) or CellTak (BD Biosciences).

### Flow cytometry

Peritoneal wash, spleen, bone marrow, mesenteric lymph node and Peyer’s Patches were collected from 8-12 week old HK2^flox^ x MB1-Cre+ mice, or Cre-littermate controls (with or without immunization). After preparation of single cell suspensions and washing, cells were counted and resuspended at 1×10^6^/mL in FACS buffer (PBS+1% FCS). Cells were stained for 30 minutes on ice with cocktails of fluorescently-labeled antibodies (see Supplementary Table 1 for details). In some experiments, cells were surface stained as above and then fixed, permeabilized and intracellularly stained. Two tubes were analyzed for every sample – one staining for HK2 as described for microscopy, and a second staining with secondary antibody alone. Cells were analyzed using a Cytoflex LX instrument equipped with 405, 488, 561 and 638nM lasers (Beckman Coulter). The resulting data were analyzed using FlowJo 10 software. For HK2 intracellular staining experiments, the mean fluorescence intensity for each population of interest was determined using both the HK2 stained and secondary-alone control tube. HK2 expression was calculated as HK2 MFI minus secondary alone background MFI for each cell population. In some experiments background subtraction was performed using population-specific background MFIs obtained from HK2 KO splenocytes stained with primary plus secondary antibody, yielding similar results.

### Mouse Immunization and Antibody Measurements

One milliliter of citrated sheep red blood cells (Colorado Serum, Denver, CO or Cedarlane Laboratories, Burlington, ON, Canada) was washed twice with 50 ml of PBS and resuspended 1:10 in PBS (0.4 ml of packed SRBCs and 3.6 ml of PBS). Two hundred microliters of the SRBC suspension were injected intraperitoneally. Flow cytometry analyses were performed at day 6 or 8 post-immunization. For antibody measurements, serum was collected on the day of immunization (day 0) and at day 5 and day10 post-immunization. Anti-SRBC Abs were measured as described^68^. Briefly, dilutions of mouse sera were incubated with washed SRBC, detected with anti-mouse IgM-PECy7 and anti-mouse IgG1-APC secondary antibodies and analyzed by flow cytometry to determine mean fluorescence intensity.

### In vitro B cell proliferation and antibody secretion assays

Splenic B cells were isolated by magnetic bead negative selection using EasySep mouse B cell isolation kit (Stem Cell Technology cat #19854). Cells were resuspended 2×10^6^ cells/mL in media containing activating factors as described above and cultured at 200uL per well in 96 well flat bottom plates for CCK8 assay or round-bottom plates for CFSE dilution and antibody secretion assays. After 3 days incubation period cell proliferation was determined using CCK8 cell viability assay, according to the manufacturer’s protocol (Sigma cat#96992). Briefly, 10uL of CCK8 reagent was added per well and after 3 hours incubation the absorbance at 450nM was measured using an ELISA plate reader. Supernatants were collected at five days for antibody measurement by ELISA assay. For cell division assays, purified B cells were labeled with CFSE prior to culture, by resuspending cells at 8×10^6^ cells per mL in PBS containing 0.6uM CFSE, incubating for 5 min at room temperature, followed by addition of an equal volume of FBS and three washes in culture medium. At the end of the culture period cells were harvested, stained with anti-IgG1-APC, anti-CD138-BV605 and analyzed by flow cytometry.

### Metabolite profiling

Metabolite profiling was performed through The Metabolomics Innovation Centre, using the Central Carbon Metabolism assay. Briefly, metabolites were extracted from snap-frozen cell pellets using 80% methanol and sonication. TCA cycle acids, sugars and aldose phosphates were quantified by LC-MRM/MS using internal standards as described^69,70^. For other metabolites, an internal standard solution containing isotope-labeled AMP, ATP, AMP, ATP, UMP, UTP, GMP, GTP, fructose-6P, fructose-bisP, UDP-glucose, glycerol-3P, NAD, NADH and glucose-1P was prepared in 50% methanol, mixed with sample, injected into a C18 column (2.1×150mm, 1.7uM) to run UPLC-MRM/MS with (-) ion detection on a Waters Acquity UPLC system coupled to a Sciex QTRAP 6500 Plus MS instrument, with the use of tributylamine acetate buffer (A) – acetonitrile-methanol (1:1) (B) as the mobile phase for gradient elution (10% to 50% B over 25min) at 0.25 mL/min and 50 degrees celcius.

### 13C-Glucose Labelling and Mass Spectrometry

Mouse splenic B-cells were stimulated (CD40L, IL-4, F(ab’)_2_ anti-IgM) overnight at 37°C in RPMI media. The following morning the cells were washed, re-plated at 37°C in Glucose free RPMI media (Gibco, Cat. #11879020) with fresh stimulation cocktail and fully labelled D-(+)-Glucose-^13^C6 (Cayman Chemical Company #26707) at a final concentration of 11.1mM, and incubated for 1 or 2 hours (glycolysis and pentose phosphate pathway analysis) or 4 hours (TCA analysis). Following labelling, cells were washed twice with cold PBS (5min x 300g at 4°C) and the cell pellets were snap frozen and stored at –80°C. Analyses of stable isotope-labeled samples was performed at the Mayo Clinic Rochester mass spectrometry core facility.

## Acknowledgements

This work was funded by the Canadian Institutes of Health Research (project grant #162268). Critical infrastructure was funded by the Canadian Foundation for Innovation. BP was funded by a studentship from the Natural Sciences and Engineering Council of Canada. EM was funded by a fellowship from Research Manitoba and CancerCare Manitoba. FA was supported by a studentship from Research Manitoba.

---

**Supplementary Figure 1.**
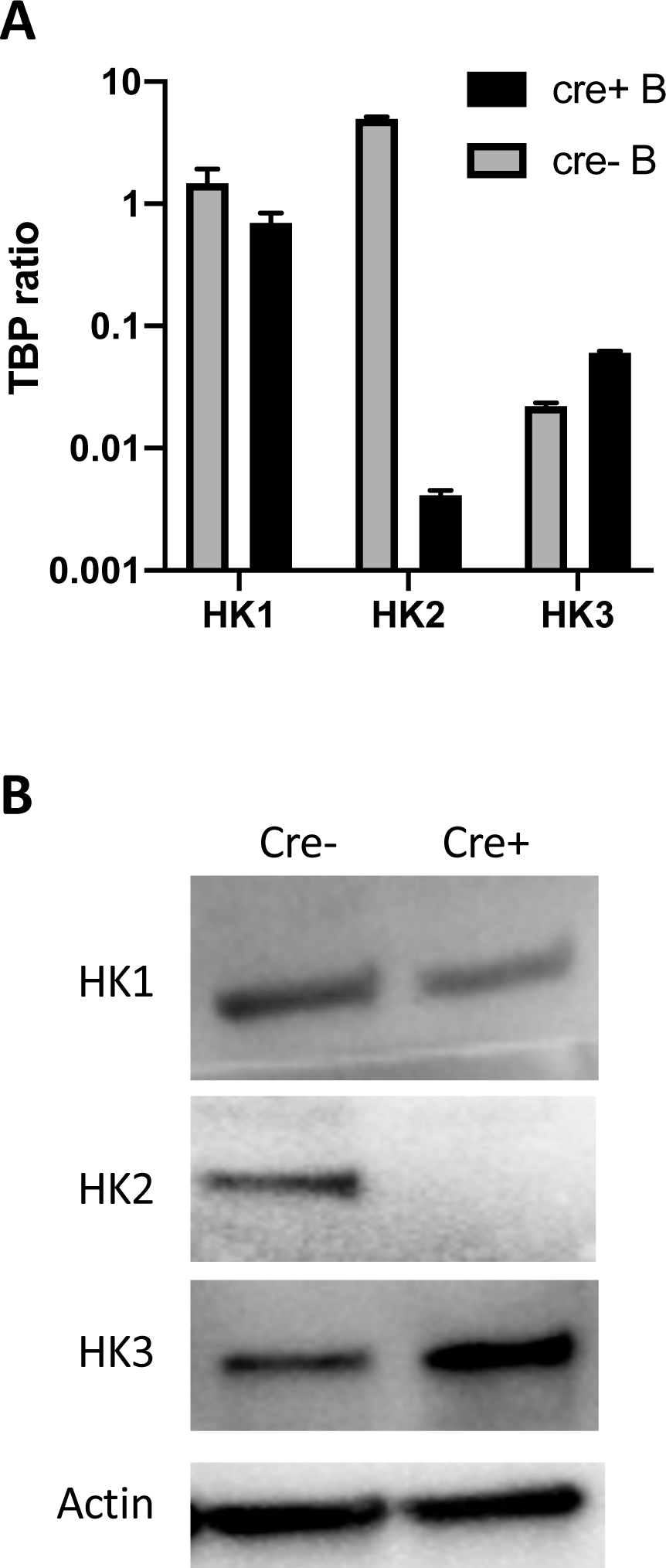
Expression of hexokinase isoforms in HK2-flox x MB1-Cre splenic B cells. B cells were isolated from spleens of HK2-flox x MB1-Cre mice or Cre negative littermates and stimulated overnight with anti-CD40+IL-4+anti-IgM. After stimulation, RNA or protein extracts were prepared for assessment of hexokinase expression. A) mRNA expression of hexokinase isoforms determined by quantitative RT-PCR assays. B) Protein expression of hexokinase isoforms determined by Western blot.

**Supplementary Figure 2.**
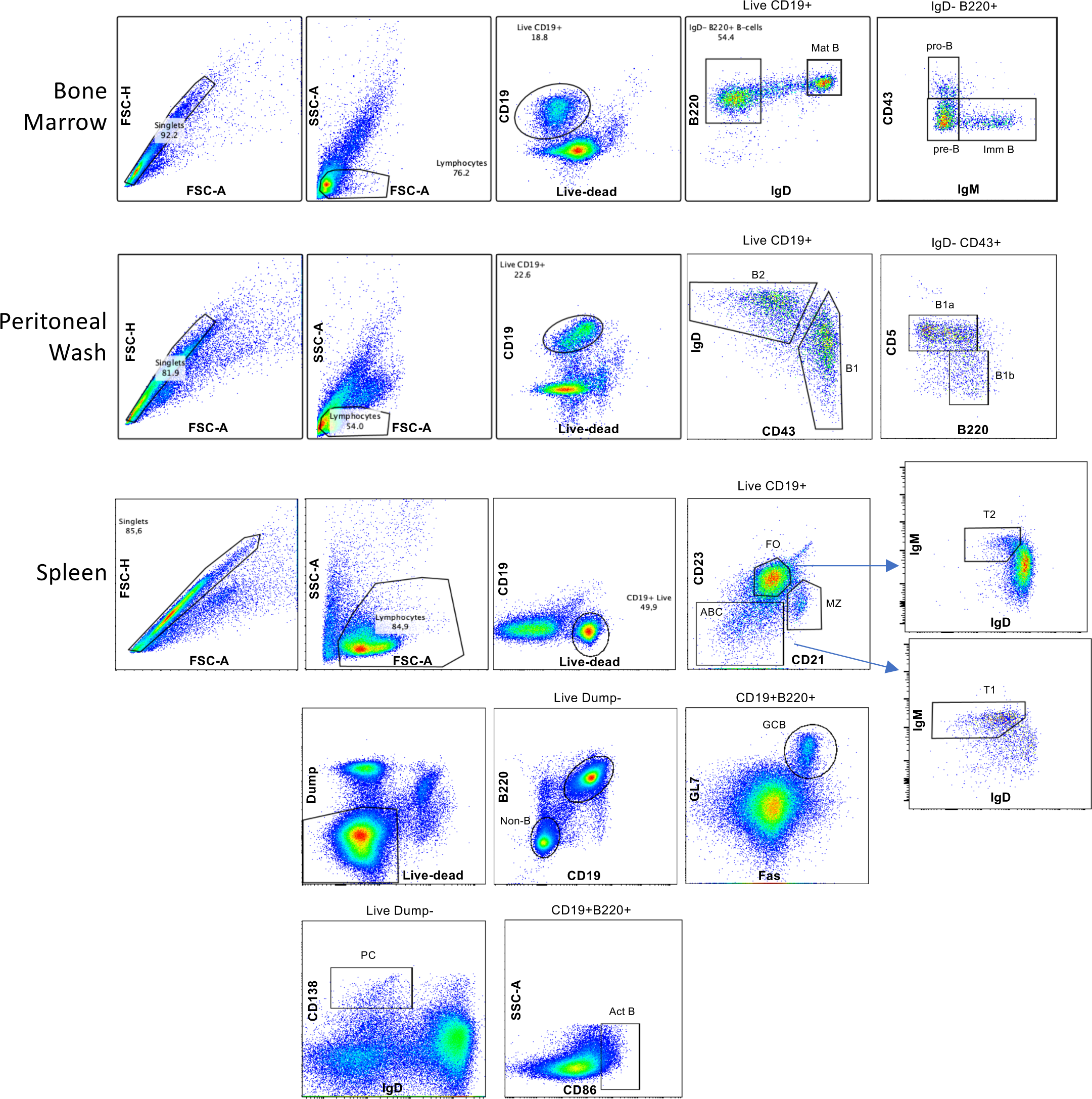
Flow gating for assessment of HK2 expression.

**Supplementary Figure 3.**
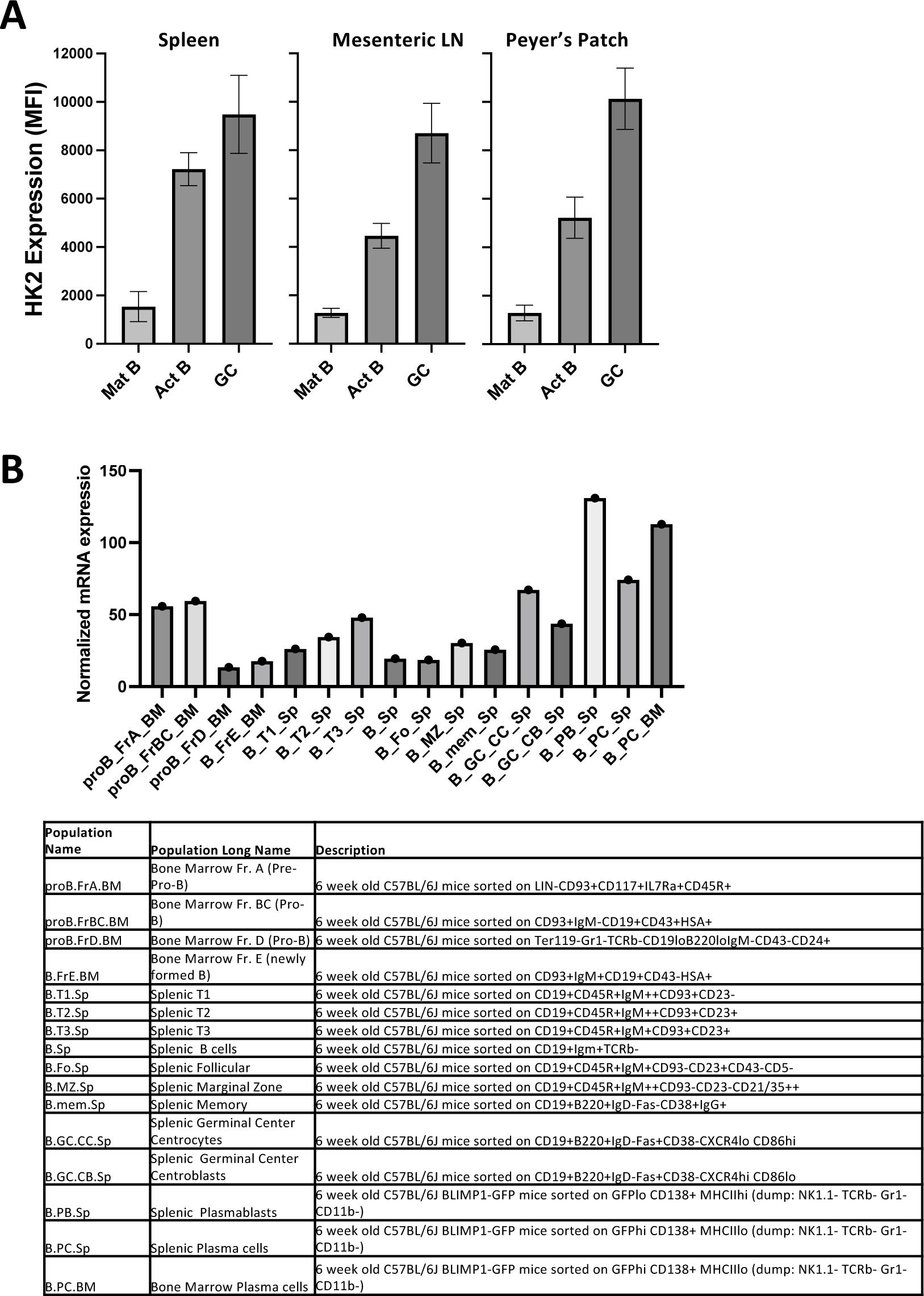
Additional HK2 expression data. A. HK2 protein expression in mature naïve B cells (CD19+CD86-IgD+; Mat B), activated B cells (CD19+CD86+IgD-; Act B) and germinal center B cells (CD19+GL7+Fas+). B. Expression of HK2 mRNA in mouse B cell subsets, from Immgen RNAseq database. Table shows the phenotypic definitions of cell populations.

**Supplementary Figure 4.**
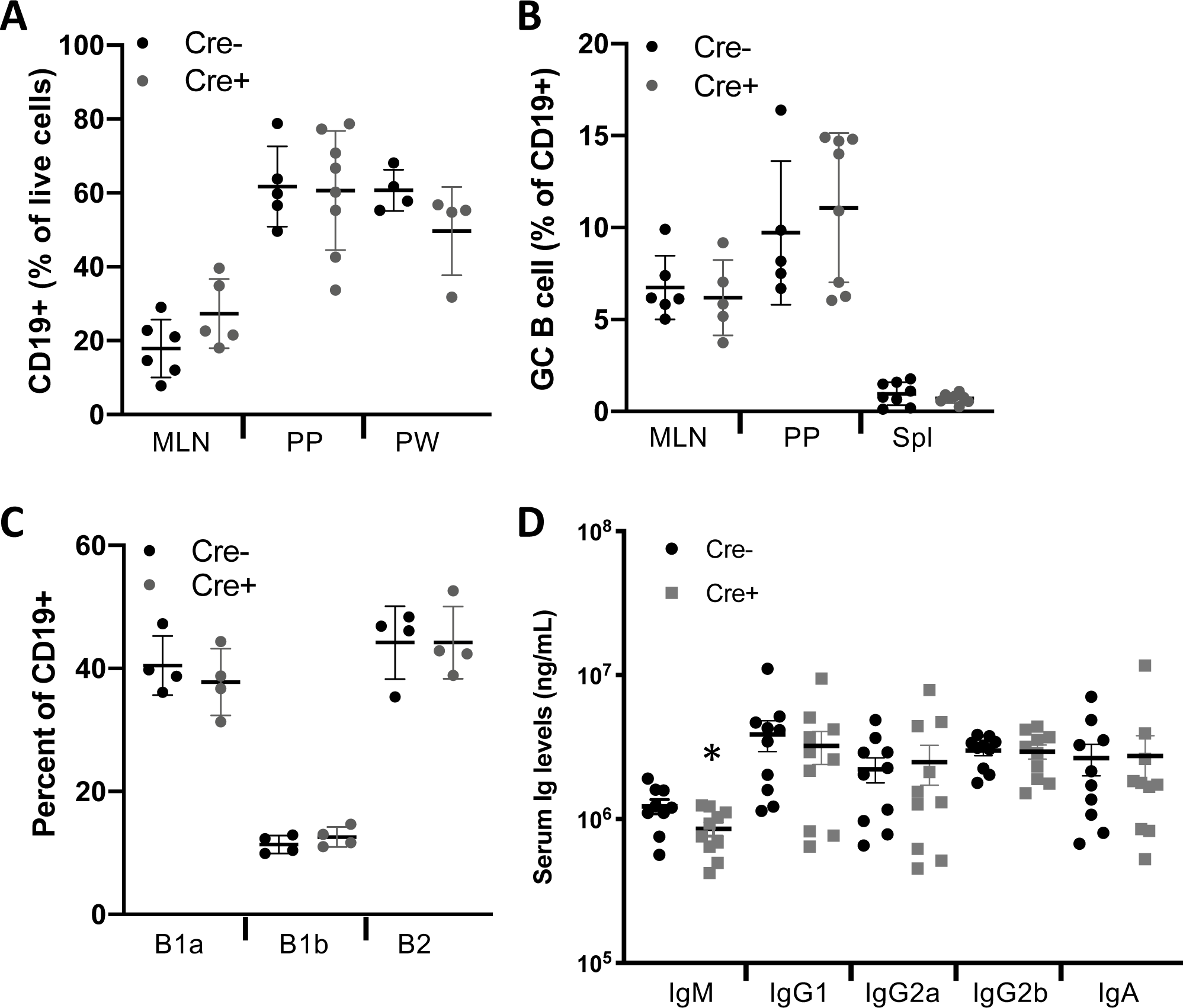
Additional characterization of HK2-deficient mice. The indicated tissues of HK2-flox x MB1-Cre mice or littermate controls were analyzed by flow cytometry to determine **A)** Total B cell frequencies, **B)** Spontaneous germinal center B cell frequencies or **C**) B1 cell subsets. BM = bone marrow, Spl = spleen, MLN = mesenteric lymph node, PP = Peyer’s patch, PW = peritoneal wash. **D)** Serum

**Supplementary Figure 5.**
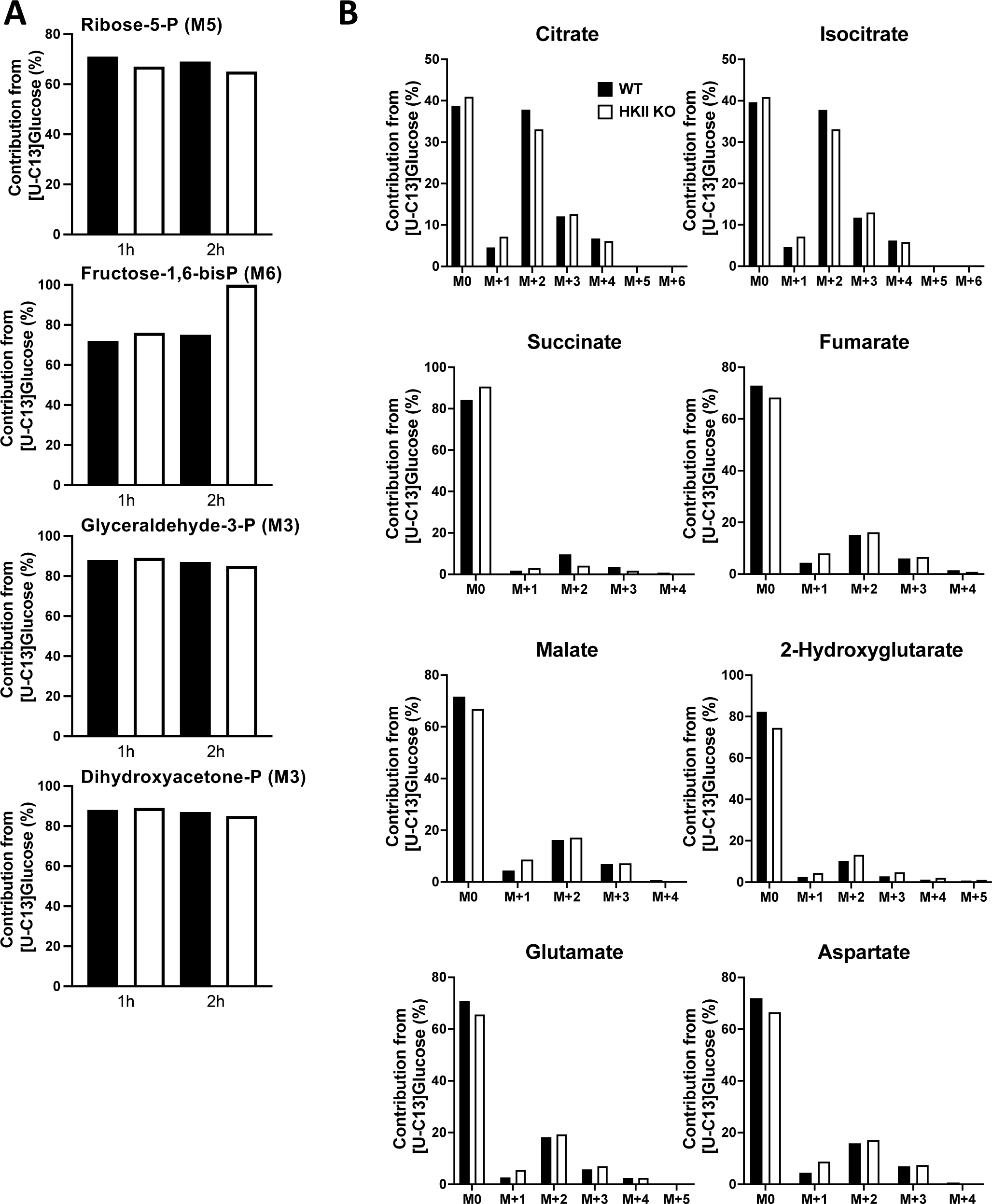
Additional metabolites assessed by 13C-labeling.

**Supplementary Figure 6.**
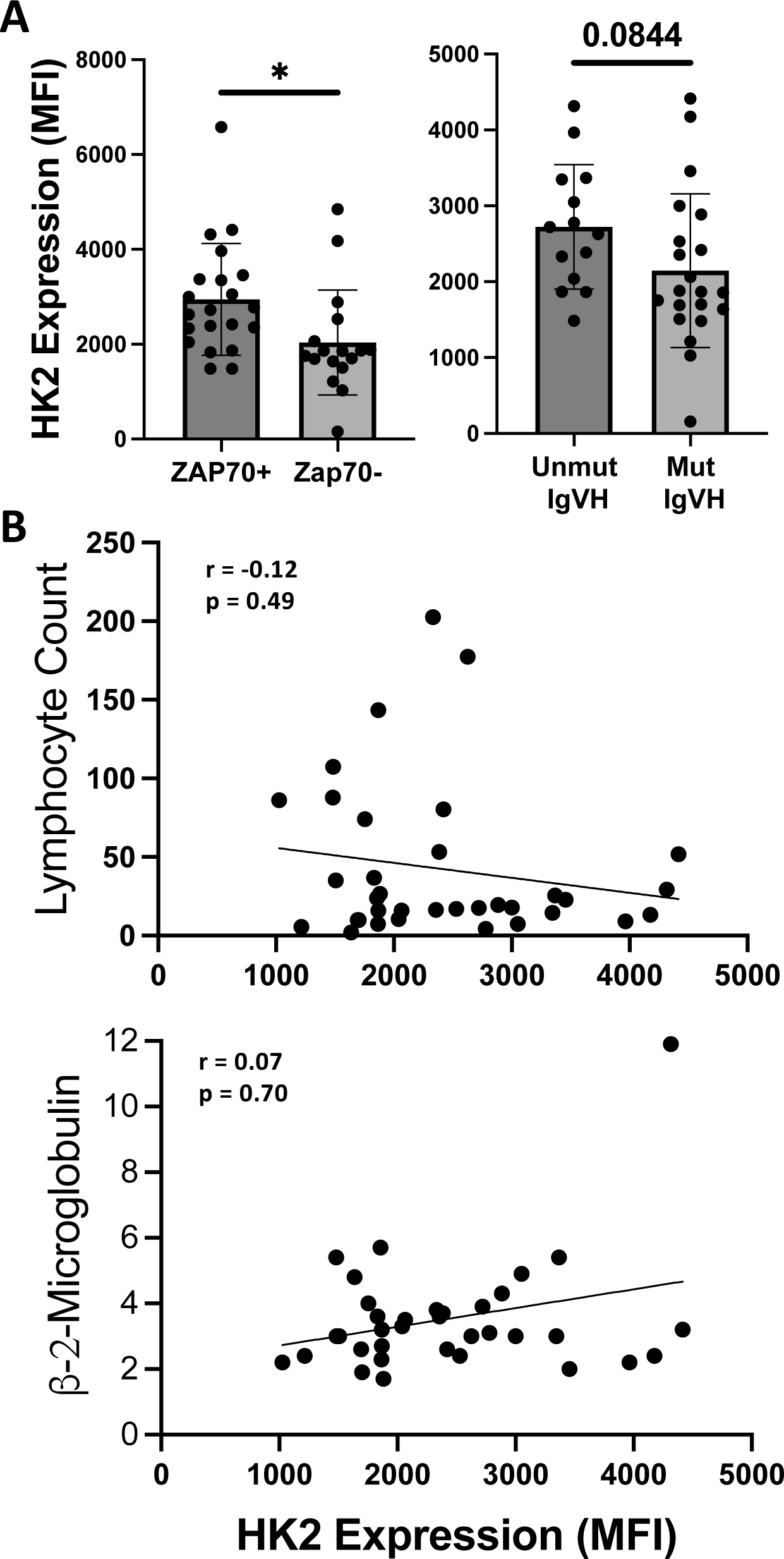
Association of HK2 expression with additional CLL prognostic and clinical parameters. A. Patients are divided into high and low risk groups based on clinical lab criteria of ZAP70 positivity (>20%) or IgVH mutation status. B. Correlation of HK2 expression with patient lymphocyte counts or beta2 microglobulin levels

Supplementary Table 1.

Supplementary Table 1
| Metabolite (nmol/mg protein) | Control Medium | HK2 KO Medium | Control STIM | HK2 KO STIM | Control STIM+2DG | HK2 KO STIM+2DG |
| --- | --- | --- | --- | --- | --- | --- |
| cAMP | 0.020044 | 0.00434 | 0.029761 | 0.035146 | 0.038025 | 0.059499 |
| c-di-AMP | 0.251122 | 0.124494 | 0.063581 | 0.077263 | 0.181087 | 0.425888 |
| AMP | 7.563542 | 2.532603 | 4.872218 | 5.875742 | 8.922186 | 14.379187 |
| ADP | 7.956866 | 3.076723 | 17.268948 | 14.938717 | 28.19063 | 34.49768 |
| ATP | 65.04215 | 15.454886 | 92.201994 | 131.394249 | 184.692931 | 334.794644 |
| CMP | 0.454213 | 0.203303 | 0.596891 | 0.470547 | 0.781073 | 1.344208 |
| cGMP | 0.001135 | 0.002056 | 0.001249 | 0.000871 | 0.003117 | 0.007829 |
| c-di-GMP | 0.000756 | 0.000457 | 0.001769 | 0.002614 | 0.000623 | 0.004697 |
| GMP | 0.35437 | 0.048427 | 0.63352 | 0.463286 | 0.721542 | 1.067851 |
| GDP | 0.879684 | 0.299243 | 1.654458 | 1.668991 | 3.757629 | 5.108303 |
| GTP | 5.384755 | 1.128672 | 5.519474 | 8.229641 | 8.902239 | 26.808999 |
| UMP | 4.818974 | 2.865197 | 2.20036 | 2.535437 | 4.57735 | 10.205645 |
| UDP | 2.628084 | 1.462409 | 3.51069 | 2.642908 | 4.246345 | 7.216603 |
| UTP | 14.194453 | 3.3915 | 44.204286 | 54.762127 | 38.933658 | 70.182992 |
| Acetyl-CoA | 12.126477 | 4.276894 | 21.380067 | 25.570763 | 26.742247 | 37.529781 |
| FAD | 0.526449 | 0.182972 | 0.52332 | 0.635384 | 0.801956 | 1.471818 |
| FMN | 0.087363 | 0.035407 | 0.075756 | 0.07247 | 0.116569 | 0.198069 |
| NAD | 8.304806 | 2.954513 | 17.941907 | 22.368282 | 21.347981 | 36.91287 |
| NADH | 6.110135 | 2.242497 | 18.043366 | 20.202021 | 18.71521 | 20.507585 |
| NADP | 0.396728 | 0.133175 | 1.343525 | 1.390003 | 0.921953 | 1.723123 |
| NADPH | 4.005474 | 1.669366 | 11.633758 | 12.093503 | 18.078133 | 23.12789 |
| Glucose | 4414.9085 | 4456.109 | 163.7025 | 481.3158 | 2023.6987 | 2718.1988 |
| Glucose 6-Phosphate | 38.1097 | 27.2647 | 13.5406 | 17.0152 | 44.1381 | 97.2057 |
| Fructose 6-Phosphate | 1.043065 | 0.603512 | 0.679203 | 1.250872 | 10.397062 | 14.381536 |
| Fructose 1,6 BisPhosphate | 1.873205 | 3.861608 | 3.229935 | 10.496692 | 4.878434 | 4.507833 |
| DiHydroxyAcetone Phosphate | 0.102491 | 0.07424 | 0.46692 | 1.404961 | 0.484664 | 0.272443 |
| Glyceraldehyde 3 Phosphate | 0.5711 | 2.8524 | 1.6197 | 3.5365 | 2.0842 | 3.5652 |
| 1,3 BisPhosphoglycerate | 0.028365 | 0.009822 | 0.007388 | 0 | 0.026493 | 0 |
| 3 PhosphoGlycerate | 3.793684 | 4.530908 | 6.195347 | 16.017345 | 7.86995 | 10.716084 |
| 2 PhosphoGlycerate | 0.166784 | 0 | 0.417803 | 0.998025 | 0.039584 | 0.805585 |
| Phosphoenolpyruvate | 0.312012 | 0.117413 | 0.585028 | 0.4598 | 0.430743 | 0.937892 |
| Pyruvate | 1309.54112 | 1103.87885 | 358.72899 | 615.77468 | 1076.84298 | 746.79081 |
| Glucose 1 Phosphate | 0.388407 | 0 | 0.01998 | 0.010747 | 0.087271 | 0.176931 |
| UDP Glucose | 2.720364 | 0.96466 | 6.942086 | 6.117261 | 11.749758 | 14.881015 |
| 6 Phosphogluconate | 0.062402 | 0.008452 | 0.195218 | 0.178488 | 0.624921 | 0.637266 |
| Ribulose 5 Phosphate | 0 | 0 | 0.40157 | 0.524718 | 3.424441 | 3.295149 |
| Ribose 5 Phosphate | 0.8093 | 2.0627 | 2.2133 | 3.2475 | 2.136 | 3.6576 |
| PRPP | 0.246962 | 0.231171 | 2.602033 | 5.047637 | 0.418588 | 2.23591 |
| Glucosamine 6 Phosphate | 0.010968 | 0 | 0 | 0 | 0 | 0.062631 |
| N-acetylglucosamine 6 phosphate | 0.102491 | 0.060077 | 0.17503 | 0.16382 | 0.078855 | 0.097077 |
| N-acetylglucosamine-1-phosphate | 0.071479 | 0.038833 | 0.647881 | 1.338446 | 0.653595 | 0.767224 |
| UDP-N-acetylglucosamine | 3.498691 | 1.137124 | 6.639894 | 6.139481 | 0.605908 | 0.776619 |
| 2 Hydroxyglutaric acid | 3.72364 | 1.71125 | 6.12918 | 3.465 | 3.00968 | 1.81117 |
| 3 Hydroxybutyric acid | 30.92233 | 12.6904 | 47.44341 | 34.12193 | 57.04519 | 9.53257 |
| Acetoacetic acid | 68.12558 | 59.68922 | 105.77044 | 118.52247 | 130.5128 | 39.73124 |
| Ethylmalonic acid | 0.73669 | 0.40272 | 0.75548 | 0.70718 | 2.12092 | 0.45173 |
| Gluconic acid | 15.35529 | 19.0181 | 31.13129 | 29.59039 | 45.09496 | 18.81706 |
| Glucuronic acid | 117.51556 | 22.88879 | 159.68437 | 27.69845 | 106.56958 | 29.81625 |
| Glutaric acid | 4.63073 | 2.47546 | 13.50549 | 6.97563 | 9.9666 | 3.87397 |
| Glyceric acid | 7.08133 | 2.73687 | 13.38258 | 5.66356 | 11.31222 | 1.61783 |
| Glycolic acid | 45.05827 | 33.37314 | 145.79605 | 83.86065 | 124.41022 | 49.22011 |
| Itaconic acid | 0.06693 | 0.06216 | 0.20668 | 0.12804 | 0.08018 | 0.10083 |
| Malonic acid | 4.6152 | 2.56419 | 5.85282 | 4.72532 | 5.6597 | 2.28069 |
| Methylmalonic acid | 0.22569 | 0 | 0.1691 | 0.33016 | 0.38122 | 0.03288 |
| Lactic acid | 3647.90236 | 1290.36821 | 2463.28781 | 2108.661 | 4241.89484 | 1236.90469 |
| Citric Acid | 42.71732 | 88.43981 | 152.59773 | 55.82248 | 61.31389 | 114.95691 |
| Aconitic acid | 0.80613 | 0.51252 | 2.76279 | 0.86564 | 1.33465 | 0.36723 |
| Isocitric acid | 0.12906 | 0.26011 | 1.26357 | 0.22567 | 1.24502 | 0.09896 |
| a-Ketoglutaric acid | 14.59782 | 30.04619 | 18.45878 | 19.31165 | 25.41704 | 26.18423 |
| Succinyl-CoA | 2.43986 | 19.948594 | 15.15972 | 27.874321 | 4.863601 | 9.475231 |
| Succinic acid | 90.28671 | 25.88925 | 112.72243 | 41.50303 | 86.11296 | 23.39468 |
| Fumaric acid | 7.23415 | 70.86022 | 51.23491 | 14.44308 | 11.94456 | 62.50916 |
| Malic acid | 22.05012 | 161.12892 | 166.51815 | 55.96843 | 31.72535 | 199.91265 |
| Oxaloacetic acid | 52.58755 | 102.27695 | 481.63036 | 98.47494 | 63.25593 | 146.81858 |

## References

1. Boothby, M. R., Brookens, S. K., Raybuck, A. L. & Cho, S. H. Supplying the trip to antibody production—nutrients, signaling, and the programming of cellular metabolism in the mature B lineage. Cell Mol Immunol 19, 352–369 (2022).

2. Jellusova, J. Metabolic control of B cell immune responses. Current Opinion in Immunology 63, 21– 28 (2020).

3. Khalsa, J. K. et al. Functionally significant metabolic differences between B and T lymphocyte lineages. Immunology 158, 104–120 (2019).

4. Jellusova, J. et al. Gsk3 is a metabolic checkpoint regulator in B cells. Nat Immunol 18, 303–312 (2017).

5. Doughty, C. A. et al. Antigen receptor–mediated changes in glucose metabolism in B lymphocytes: role of phosphatidylinositol 3-kinase signaling in the glycolytic control of growth. Blood 107, 4458– 4465 (2006).

6. Jellusova, J. & Rickert, R. C. The PI3K pathway in B cell metabolism. Critical Reviews in Biochemistry and Molecular Biology 51, 359–378 (2016).

7. Jellusova, J. Metabolic control of B cell immune responses. Current Opinion in Immunology 63, 21– 28 (2020).

8. Jayachandran, N. et al. TAPP Adaptors Control B Cell Metabolism by Modulating the Phosphatidylinositol 3-Kinase Signaling Pathway: A Novel Regulatory Circuit Preventing Autoimmunity. J Immunol 201, 406–416 (2018).

9. Akkaya, M. et al. Second signals rescue B cells from activation-induced mitochondrial dysfunction and death. Nat Immunol 19, 871–884 (2018).

10. Waters, L. R., Ahsan, F. M., Wolf, D. M., Shirihai, O. & Teitell, M. A. Initial B Cell Activation Induces Metabolic Reprogramming and Mitochondrial Remodeling. iScience 5, 99–109 (2018).

11. Caro-Maldonado, A. et al. Metabolic reprogramming is required for antibody production that is suppressed in anergic but exaggerated in chronically BAFF-exposed B cells. J Immunol 192, 3626– 3636 (2014).

12. Patke, A., Mecklenbrauker, I., Erdjument-Bromage, H., Tempst, P. & Tarakhovsky, A. BAFF controls B cell metabolic fitness through a PKC beta– and Akt-dependent mechanism. The Journal of experimental medicine 203, 2551–2562 (2006).

13. Abbott, R. K. et al. Germinal Center Hypoxia Potentiates Immunoglobulin Class Switch Recombination. J Immunol 197, 4014–4020 (2016).

14. Cho, S. H. et al. Germinal centre hypoxia and regulation of antibody qualities by a hypoxia response system. Nature 537, 234–238 (2016).

15. Raybuck, A. L. et al. B Cell-Intrinsic mTORC1 Promotes Germinal Center-Defining Transcription Factor Gene Expression, Somatic Hypermutation, and Memory B Cell Generation in Humoral Immunity. J Immunol 200, 2627–2639 (2018).

16. Dominguez-Sola, D. et al. The proto-oncogene MYC is required for selection in the germinal center and cyclic reentry. Nat Immunol 13, 1083–1091 (2012).

17. Ersching, J. et al. Germinal Center Selection and Affinity Maturation Require Dynamic Regulation of mTORC1 Kinase. Immunity 46, 1045–1058.e6 (2017).

18. Lam, W. Y. et al. Mitochondrial pyruvate import promotes long-term survival of antibody-secreting plasma cells. Immunity 45, 60–73 (2016).

19. Corcoran, L. M. & Nutt, S. L. Long-Lived Plasma Cells Have a Sweet Tooth. Immunity 45, 3–5 (2016).

20. Roberts, D. J. & Miyamoto, S. Hexokinase II integrates energy metabolism and cellular protection: Akting on mitochondria and TORCing to autophagy. Cell Death Differ 22, 248–257 (2015).

21. Okkenhaug, K. Signaling by the phosphoinositide 3-kinase family in immune cells. Annu Rev Immunol 31, 675–704 (2013).

22. Fruman, D. A. et al. The PI3K Pathway in Human Disease. Cell 170, 605–635 (2017).

23. Doughty, C. A. et al. Antigen receptor-mediated changes in glucose metabolism in B lymphocytes: role of phosphatidylinositol 3-kinase signaling in the glycolytic control of growth. Blood 107, 4458– 4465 (2006).

24. Jellusova, J. et al. Gsk3 is a metabolic checkpoint regulator in B cells. Nat Immunol 18, 303–312 (2017).

25. Fruman, D. A. Phosphoinositide 3-kinase and its targets in B-cell and T-cell signaling. Current opinion in immunology 16, 314–320 (2004).

26. Balla, T. Phosphoinositides: tiny lipids with giant impact on cell regulation. Physiol Rev 93, 1019–137 (2013).

27. Lemmon, M. A. Pleckstrin homology (PH) domains and phosphoinositides. Biochemical Society symposium (74), 81–93 (2007).

28. Fruman, D. & Limon, J. Akt and mTOR in B Cell Activation and Differentiation. Frontiers in Immunology 3, (2012).

29. Memmott, R. M. & Dennis, P. A. Akt-dependent and independent mechanisms of mTOR regulation in cancer. Cell Signal 21, 656–664 (2009).

30. Zhu, Z., Shukla, A., Ramezani-Rad, P., Apgar, J. R. & Rickert, R. C. The AKT isoforms 1 and 2 drive B cell fate decisions during the germinal center response. Life Science Alliance 2, (2019).

31. Dufort, F. J. et al. Cutting Edge: IL-4-Mediated Protection of Primary B Lymphocytes from Apoptosis via Stat6-Dependent Regulation of Glycolytic Metabolism. The Journal of Immunology 179, 4953– 4957 (2007).

32. Maratou, E. et al. Glucose transporter expression on the plasma membrane of resting and activated white blood cells. Eur J Clin Invest 37, 282–290 (2007).

33. Wieman, H. L., Wofford, J. A. & Rathmell, J. C. Cytokine stimulation promotes glucose uptake via phosphatidylinositol-3 kinase/Akt regulation of Glut1 activity and trafficking. Mol Biol Cell 18, 1437– 1446 (2007).

34. Nawaz, M. H., et al. The catalytic inactivation of the N-half of human hexokinase 2 and structural and biochemical characterization of its mitochondrial conformation. Bioscience Reports 38, BSR20171666 (2018).

35. Robey, R. B. & Hay, N. Mitochondrial hexokinases, novel mediators of the antiapoptotic effects of growth factors and Akt. Oncogene 25, 4683–4696 (2006).

36. Rodríguez-Saavedra, C. et al. Moonlighting Proteins: The Case of the Hexokinases. Front. Mol. Biosci. 8, 701975 (2021).

37. Varanasi, S. K., Jaggi, U., Hay, N. & Rouse, B. T. Hexokinase II may be dispensable for CD4 T cell responses against a virus infection. PLoS One 13, e0191533 (2018).

38. De Jesus, A. et al. Hexokinase 1 cellular localization regulates the metabolic fate of glucose. Molecular Cell 82, 1261–1277.e9 (2022).

39. Roberts, D. J., Tan-Sah, V. P., Smith, J. M. & Miyamoto, S. Akt Phosphorylates HK-II at Thr-473 and Increases Mitochondrial HK-II Association to Protect Cardiomyocytes. Journal of Biological Chemistry 288, 23798–23806 (2013).

40. Wray-Dutra, M. N. et al. Activated PIK3CD drives innate B cell expansion yet limits B cell-intrinsic immune responses. J Exp Med 215, 2485–2496 (2018).

41. Hoellenriegel, J. et al. The phosphoinositide 3’-kinase delta inhibitor, CAL-101, inhibits B-cell receptor signaling and chemokine networks in chronic lymphocytic leukemia. Blood 118, 3603–12 (2011).

42. Vangapandu, H. V. et al. B-cell Receptor Signaling Regulates Metabolism in Chronic Lymphocytic Leukemia. Molecular Cancer Research 15, 1692–1703 (2017).

43. Roy Chowdhury, S., et al. Mitochondrial Respiration Correlates with Prognostic Markers in Chronic Lymphocytic Leukemia and Is Normalized by Ibrutinib Treatment. Cancers 12, 650 (2020).

44. Calissano, C. et al. Intraclonal Complexity in Chronic Lymphocytic Leukemia: Fractions Enriched in Recently Born/Divided and Older/Quiescent Cells. Mol Med 17, 1374–1382 (2011).

45. Herndon, T. M. et al. Direct in vivo evidence for increased proliferation of CLL cells in lymph nodes compared to bone marrow and peripheral blood. Leukemia 31, 1340–1347 (2017).

46. Jacobs, S. R. et al. Glucose Uptake Is Limiting in T Cell Activation and Requires CD28-Mediated Akt-Dependent and Independent Pathways1. The Journal of Immunology 180, 4476–4486 (2008).

47. Rathmell, J. C. et al. Akt-Directed Glucose Metabolism Can Prevent Bax Conformation Change and Promote Growth Factor-Independent Survival. Molecular and Cellular Biology 23, 7315–7328 (2003).

48. Siska, P. J. et al. Suppression of Glut1 and Glucose Metabolism by Decreased Akt/mTORC1 Signaling Drives T Cell Impairment in B Cell Leukemia. The Journal of Immunology 197, 2532–2540 (2016).

49. Dibble, C. C. & Cantley, L. C. Regulation of mTORC1 by PI3K signaling. Trends Cell Biol 25, 545–555 (2015).

50. Saxton, R. A. & Sabatini, D. M. mTOR Signaling in Growth, Metabolism, and Disease. Cell 168, 960– 976 (2017).

51. Zeng, H. & Chi, H. mTOR and lymphocyte metabolism. Curr Opin Immunol 25, 347–355 (2013).

52. Kim, J., Gao, P., Liu, Y.-C., Semenza, G. L. & Dang, C. V. Hypoxia-inducible factor 1 and dysregulated c-Myc cooperatively induce vascular endothelial growth factor and metabolic switches hexokinase 2 and pyruvate dehydrogenase kinase 1. Mol Cell Biol 27, 7381–7393 (2007).

53. Broecker-Preuss, M., Becher-Boveleth, N., Bockisch, A., Dührsen, U. & Müller, S. Regulation of glucose uptake in lymphoma cell lines by c-MYC– and PI3K-dependent signaling pathways and impact of glycolytic pathways on cell viability. J Transl Med 15, 158 (2017).

54. Dong, Y., Tu, R., Liu, H. & Qing, G. Regulation of cancer cell metabolism: oncogenic MYC in the driver’s seat. Sig Transduct Target Ther 5, 1–11 (2020).

55. Wilson, J. E. Isozymes of mammalian hexokinase: structure, subcellular localization and metabolic function. Journal of Experimental Biology 206, 2049–2057 (2003).

56. Roberts, D. J. & Miyamoto, S. Hexokinase II integrates energy metabolism and cellular protection: Akting on mitochondria and TORCing to autophagy. Cell Death Differ 22, 248–257 (2015).

57. Yang, T. et al. PIM2-mediated phosphorylation of hexokinase 2 is critical for tumor growth and paclitaxel resistance in breast cancer. Oncogene 37, 5997–6009 (2018).

58. Ciscato, F. et al. Hexokinase 2 displacement from mitochondria-associated membranes prompts Ca^2+^ –dependent death of cancer cells. EMBO Reports 21, e49117 (2020).

59. Brookens, S. K. et al. Plasma Cell Differentiation, Antibody Quality, and Initial Germinal Center B Cell Population Depend on Glucose Influx Rate. J Immunol 212, 43–56 (2024).

60. Tsai, S. et al. Insulin Receptor-Mediated Stimulation Boosts T Cell Immunity during Inflammation and Infection. Cell Metabolism 28, 922–934.e4 (2018).

61. Tabatabaei Shafiei, M., et al. Detecting glycogen in peripheral blood mononuclear cells with periodic acid schiff staining. J Vis Exp 52199 (2014) doi:10.3791/52199.

62. Chen, D. et al. Coupled analysis of transcriptome and BCR mutations reveals role of OXPHOS in affinity maturation. Nat Immunol 22, 904–913 (2021).

63. Sharma, R. et al. Distinct metabolic requirements regulate B cell activation and germinal center responses. Nat Immunol 24, 1358–1369 (2023).

64. Rozovski, U., Hazan-Halevy, I., Barzilai, M., Keating, M. J. & Estrov, Z. Metabolism pathways in chronic lymphocytic leukemia. Leuk Lymphoma 57, 758–765 (2016).

65. Meadows, S. A. et al. PI3Kdelta inhibitor, GS-1101(CAL-101), attenuates pathway signaling, induces apoptosis, and overcomes signals from the microenvironment in cellular models of Hodgkin lymphoma. Blood.

66. Choi, S.-C. et al. Inhibition of glucose metabolism selectively targets autoreactive follicular helper T cells. Nat Commun 9, 4369 (2018).

67. Patra, K. C. et al. Hexokinase 2 is required for tumor initiation and maintenance and its systemic deletion is therapeutic in mouse models of cancer. Cancer Cell 24, 213–228 (2013).

68. McAllister, E. J., Apgar, J. R., Leung, C. R., Rickert, R. C. & Jellusova, J. New methods to analyze B cell immune responses to the thymus dependent antigen sheep red blood cells. J Immunol 199, 2998– 3003 (2017).

69. Han, J., Gagnon, S., Eckle, T. & Borchers, C. H. Metabolomic Analysis of Key Central Carbon Metabolism Carboxylic Acids as Their 3-Nitrophenylhydrazones by UPLC/ESI-MS. Electrophoresis 34, 2891–2900 (2013).

70. Han, J., Tschernutter, V., Yang, J., Eckle, T. & Borchers, C. H. Analysis of selected sugars and sugar phosphates in mouse heart tissue by reductive amination and liquid chromatography-electrospray ionization mass spectrometry. Anal Chem 85, 5965–5973 (2013).

